# Small viruses reveal bidirectional evolution between HK97-fold viruses and encapsulins via procapsids

**DOI:** 10.1101/2025.06.18.659913

**Authors:** Abelardo Aguilar, Ingrid Miranda, Omer Nadel, Stephen Nayfach, Luis Ruiz Pestana, Anca M. Segall, Forest L. Rohwer, Simon Roux, Simon White, Antoni Luque

## Abstract

The HK97-fold is an ancient, conserved protein structure that forms protein shells. It is the building block of capsids that protect the genome of viruses in the *Duplodnaviria* realm, infecting organisms across all domains of life. It is also the building block of encapsulins, compartments that confine biochemical reactions in prokaryotes. Recent genomic studies have hypothesized that encapsulins evolved from HK97-fold viruses. However, this evolutionary pathway is challenging to justify biophysically because HK97-fold viruses form larger and more complex protein shells than encapsulins. We addressed this paradox by searching for smaller and simpler HK97-fold viral capsids across ecosystems. The investigation yielded a well-defined group of viral entities, which encode HK97-fold proteins displaying molecular similarities with encapsulins and lacking portal and tail genes. The structural phylogenetic analysis of HK97-fold entities revealed bidirectional evolutionary transitions between encapsulins and HK97-fold viruses. An evolutionary mechanism responsible for such transitions was proposed based on the presence of lysogeny-associated genes in the viral entities and the structural and molecular parallels between encapsulins and procapsids, the immature state of viral capsids before genome packaging. We concluded that procapsids are akin to the common HK97-fold protein shell ancestor and might still facilitate transitions between modern viruses and encapsulins. The potential genomic and biochemical functions of HK97-fold procapsids make them an ideal candidate model for the early evolution of life and might require a revision of the role of viral capsids in the biosphere.

## INTRODUCTION

One longstanding debate on the origin of life addresses whether life began primarily through the emergence of self-replicating informational molecules endowed with catalytic activity (a genetics-first model), like the RNA world hypothesis (Gilbert, 1986; Pressman et al., 2015), or as a cooperative network of autocatalytic reactions (a metabolism-first model) that preceded genetic polymers (Kauffman, 1971; Smith & Morowitz, 2004; Takagi et al., 2020; Wächtershäuser, 1997). Since modern cells require both metabolism and genetics, recent formulations on the origin of life explore hybrid scenarios in which primordial replicators and metabolic systems co-sevolved and became functionally integrated (Becerra et al., 2007; Preiner et al., 2020; Vincent et al., 2019). This view acknowledges that informational and metabolic processes likely became linked early on, rather than one strictly preceding the other. A critical step in any origin-of-life scenario is the emergence of compartments that could concentrate reactants, protect fragile molecules, and favor the exchange of molecules (Cornell et al., 2019; Mansy & Szostak, 2009; Segré et al., 2001). Indeed, life at its core is about creating distinct chemical spaces through compartmentalization. Contemporary biological compartments can be interpreted as imprints or milestones in the evolution of life and are invaluable for extracting clues on the origin of life (Caetano-Anollés et al., 2009). Despite the central importance of compartments, one ancient group of compartment-forming proteins (Duda & Teschke, 2019; Krupovic & Koonin, 2017), which form icosahedral shells capable of performing both genomic and metabolic functions (Giessen, 2016; Suhanovsky & Teschke, 2015; Sutter et al., 2008; Wikoff et al., 2000) and are one of the most frequent proteins on the planet (Cobián Güemes et al., 2016; Koonin et al., 2024) has been overlooked in the debate about the origin of life.

This highly diverse group of proteins in sequence space (Lee et al., 2022) is characterized by displaying the HK97-fold or Johnson fold (Hendrix, 2005; Wikoff et al., 2006). This fold specializes in assembling icosahedral shells (Figure 1A). The subset of HK97-fold proteins with genomic functions forms the capsids protecting the genome of viruses in the *Duplodnaviria* realm (Koonin et al., 2020; Krupovic & Koonin, 2017). This includes tailed prokaryotic viruses (or tailed phages) that infect bacteria and archaea (Hua et al., 2017; Pietilä et al., 2013; Suhanovsky & Teschke, 2015; Wikoff et al., 2000), related prokaryotic mobile elements, like phage inducible chromosomal islands (PICIs) and gene transfer agents (GTAs) (Bárdy et al., 2020; Dearborn et al., 2017) and herpesviruses infecting animals (Dai & Hong Zhou, 2018; Heymann et al., 2003). The other subset of HK97-fold proteins forms encapsulins, cellular compartments carrying ancient metabolic functions in bacteria and archaea, including iron storage (Giessen et al., 2019), sulfur metabolism (Nichols et al., 2021), or oxidative stress response (Jones et al., 2024; Sutter et al., 2008). The current evolutionary paradigm of HK97-fold proteins, based on structural and phylogenetic analyses and presence across taxa, indicates that encapsulins are derived from HK97-fold viruses. However, this paradigm does not provide a rationale for how HK97-fold shells evolved metabolic functions from shells carrying genomic functions and how HK97-fold viruses, which form far more complex structures than encapsulins (Luque et al., 2020), would have emerged before encapsulins. The differences in structural complexity of HK97-fold compartments were not considered in prior evolutionary studies, but, as we show here, they are key to elucidating the common ancestor of HK97-fold compartments and the function it may have carried in early life.

**Figure 1.**
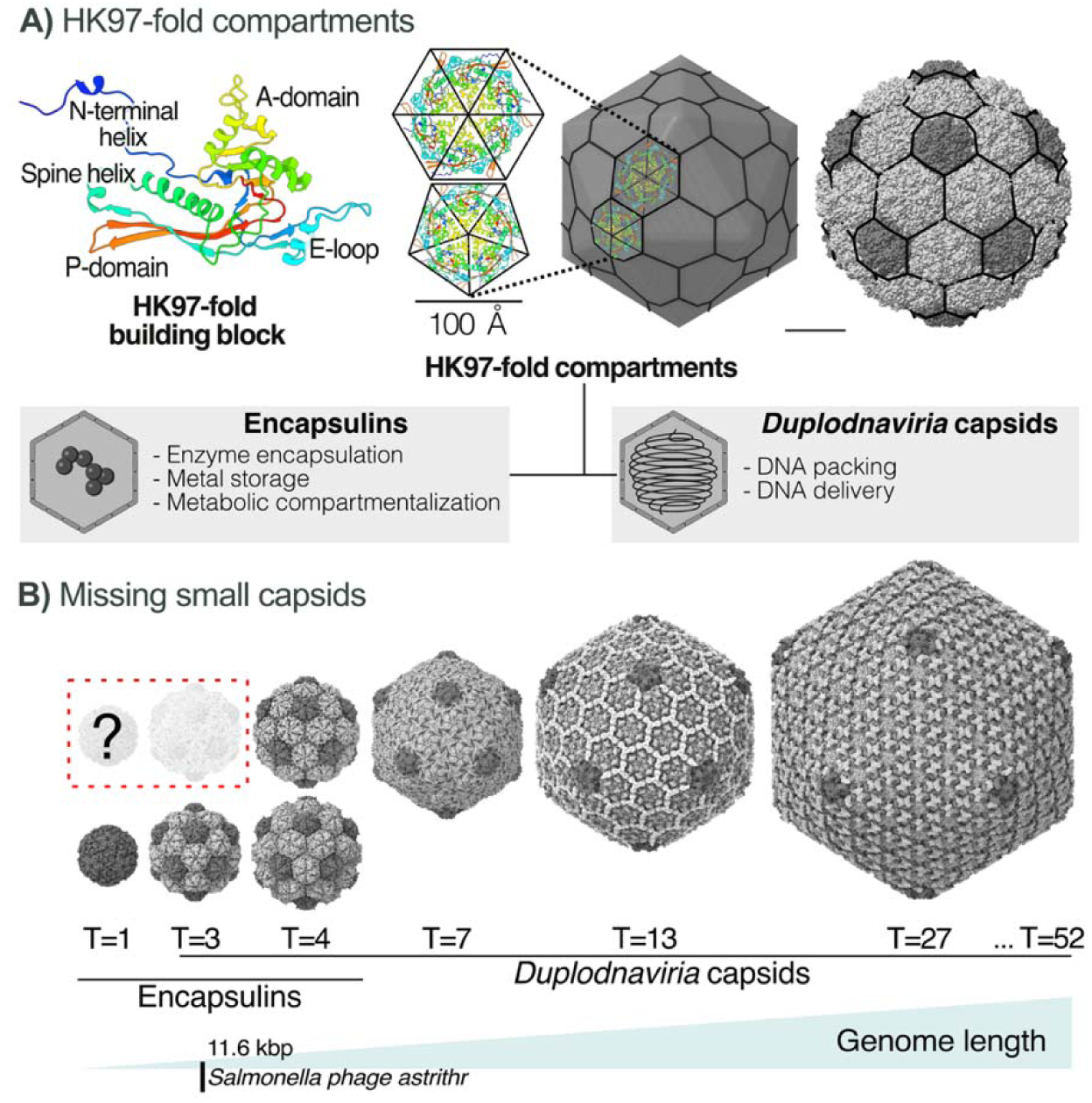
Size distribution of HK97-fold biochemical and genomic compartments. A) Schematic representatio of the HK97-fold and its main domains displayed with different colors. The geometrical and rendered capsi illustrates the distribution of capsomers as pentamers (five proteins) and hexamers (six proteins). The gray boxes describe the two main compartments. B) Size range of HK97-fold compartments showcasing geometric capsi architectures (T-numbers). The dashed red box indicates missing (question mark) and non-reconstructed small capsids among *Duplodnaviria*. The blue scale gradient indicates the genome size gradient. The vertical line indicates the smallest *Duplodnaviria* genome as a reference scale.

The HK97-fold proteins in encapsulins co-assemble around several copies of a protein called cargo, which is encoded in the same operon and is responsible for catalyzing the biochemical reactions (Chmelyuk et al., 2023; Jones et al., 2024). In some encapsulins, cargo proteins favor the formation of larger icosahedral capsids (Gómez-Barrera et al., 2025). Most encapsulins form protein shells built from 60 HK97-fold proteins distributed in 12 pentamer (oligomers of five proteins) as the vertices of the simplest icosahedral architecture, T=1, and several form T=3 and a few T=4 icosahedral architectures combining pentamers at the vertice and hexamers on the faces (Andreas & Giessen, 2021; Chmelyuk et al., 2023; Kashif-Khan et al., 2024) (Figure 1B). The T-number (or triangulation number) is the number of major capsid proteins in the asymmetric unit, which follows a sequence (T=1, 3, 4, 7,…) defined by the equation T = h^2^ + h·k + k^2^, and determines the number of major capsid proteins, 60T, in an icosahedral shell (Caspar & Klug, 1962; Twarock & Luque, 2019).

HK97-fold viruses instead undergo a complex maturation process requiring scaffold proteins to guide the assembly of the correct capsid size, followed often by the assembly of minor proteins reinforcing the structure (Dearborn et al., 2017; Fokine & Rossmann, 2014; Hardy et al., 2020; Morais et al., 2003; Podgorski et al., 2023). The HK97-fold proteins guided by the scaffold proteins assemble around a portal complex, which is responsible for the translocation of the double-stranded (ds) DNA genome in the capsid (Chen et al., 2011; Ibarra et al., 2000; Ignatiou et al., 2019; Woodson et al., 2021). This forms an empty shell called a procapsid, which, as the dsDNA is packaged through the portal, expands, releases the scaffold proteins, and becomes more faceted (Ionel et al., 2011; Oh et al., 2014; Roos et al., 2012; Veesler & Johnson, 2012; Wikoff et al., 2006). Prokaryotic HK97-fold viruses and mobile elements append to the portal a protein tail that enables the delivery of dsDNA to the host (Hawkins et al., 2023; Rao et al., 2021; Xu et al., 2019). HK97-fold viral genomes display a wide range of lengths due to the versatility of HK97-fold proteins forming capsid architectures of different sizes (Berg & Roux, 2021; Hua et al., 2017; Luque et al., 2020; Suhanovsky & Teschke, 2015). Most HK97-type viruses form primarily T=7 structures followed by T=13 and T=16 structures (Dai & Hong Zhou, 2018; Helgstrand et al., 2003; Lander et al., 2013; Luque et al., 2020; Suhanovsky & Teschke, 2015; Yap & Rossmann, 2014) (Figure 1B). The largest HK97-fold viral capsid reconstructed corresponds to Lysinbacillus Phage G, forming a T=52 architecture, containing more than 3,000 major capsid proteins and packaging a 626 kb-long dsDNA genome (González et al., 2020; Hua et al., 2017). The smallest reconstructions correspond to elongated T=3 capsids of Bacillus phage phi29 (Choi et al., 2006) and quasi-spherical T=4 capsids of *Streptococci C1 phage* (Aksyuk et al., 2012) the satellite enterobacteria phage *P4* (Kizziah et al., 2020), and the *Staphylococcus aureus* pathogenic island 1 or SaPI1 (Dearborn et al., 2017). The smallest tailed phage genomes are 11.6 kb-long from the non-satellite *Salmonella phage Astrid* (Olsen et al., 2020) and the satellite phage P4 (Haggård-Liungquist et al., 1995). The smallest HK97-fold capsid characterized, associated with genomic functions, corresponds to a *Rhodobacter capsulatus* gene transfer agent (RcGTA), encoded by a 15 kb-long genomic region that assembles a shortened oblate-shaped T=3-like structure that packages 4.0–4.5 kb long dsDNA host fragments (Bárdy et al., 2020). No isolated HK97-fold viruses or mobile elements have been described to form the simplest icosahedral architecture, T=1 (Luque et al., 2020).

The structural complexity of HK97-fold compartments, combined with the principle of parsimony, suggests that HK97-fold viruses evolved from encapsulins, challenging the current paradigm. To resolve this contradiction, we hypothesized that HK97-fold compartments evolved from simpler HK97-fold viruses. We built our approach on three key elements. First, genomes associated with HK97-fold viruses, in particular tailed-like phages, are prevalent when sequencing environmental samples and display a much higher diversity than their isolated counterparts (Cobián Güemes et al., 2016; Koonin et al., 2024; Paez-Espino et al., 2016; Wigington et al., 2016; Wommack & Colwell, 2000). Second, the genome length of HK97-fold viruses follows a well-established allometric relationship with their capsid size and architecture (Cui et al., 2014; Lee et al., 2022; Luque et al., 2020). Third, tailed phages can encode direct terminal repeats (DTRs) in their genomes, providing a bioinformatic proxy for identifying complete genomes among uncultured viruses (Benler et al., 2021; Casjens & Gilcrease, 2009; Nayfach et al., 2021). A previous study combined these elements to search for simple HK97-fold viral genomes in human gut metagenomes (Luque et al., 2020). It found 14 candidates encoding HK97-fold proteins in genomes below the isolated threshold of 11.6 kb, which we defined here as the twilight zone of HK97-fold viral genomes. The genomes ranged from 6.9 kb to 10.0 kb, with no candidates in the range of 1 kb to 5 kb predicted to be compatible with T=1 architectures.

Here, we expanded the analysis to include samples across ecosystems, refined the HK97-fold detection strategies, introduced a new network method comparing HK97-fold profiles, and generated a large-scale structural phylogenetic analysis compared to previous studies. Our investigation identified 523 HK97-fold-encoding genomes across the twilight zone, including 93 encoding HK97-fold proteins molecularly related to encapsulins. The genomes lacked portal and tail genes, and their HK97-fold proteins revealed at least four instances of evolution between encapsulins and HK97-fold viruses, suggesting bidirectional evolution. Further molecular and structural comparisons led us to propose that procapsids are the evolutionary nexus and extant common ancestor of HK97-fold compartments. We concluded by discussing how procapsids could carried both genomic and biochemical functions in early life and modern viral infections.

## METHODS

The links to access the Data Files referenced in the Methods and Results sections are available in the Supplementary Materials file. Code and analysis workflows related to this study are available at https://github.com/luquelab/virus_encapsulin_2025.

### Identifying environmental HK97-fold viral genomes in the twilight zone

Small, complete HK97-fold viral genomes were identified from viral contigs containing direct terminal repeats (DTRs) in public environmental omics data (metagenomes, metatranscriptomes, and metaviromes) used in the viral pipeline CheckV (Nayfach et al., 2021). This included over 14.4 billion contigs from IMG/M (I.-M. A. Chen et al., 2019), Mgnify (Mitchell et al., 2019), human microbiomes (Nayfach et al., 2019; Pasolli et al., 2019; Soto-Perez et al., 2019), and ocean virome (Paez-Espino et al., 2019). The complete list of IMG datasets used in this study i provided in Data File S1. The standard hierarchical metagenomic biome classification schema was used (Ivanova et al., 2010), available in Data File S2. The assignment of contigs to biomes i provided in Data File S3.

The bioinformatics pipeline CheckV estimated the completeness of viral genomes and removed non-viral regions from integrated proviruses (Nayfach et al., 2021). The viral contigs selected for analysis (Data File S4) satisfied three criteria. First, the contigs contained a direct terminal repeat (DTR) window of at least 20 base pairs (bp), which was used as a bioinformatic proxy for a complete or closed viral genome (Benler et al., 2021; Casjens & Gilcrease, 2009; Nayfach et al., 2021). Second, each selected genome contained at least one marker gene associated with tailed phages, including major capsid protein, portal protein, terminase, baseplate, or tail genes (Koonin et al., 2020). Open reading frames were identified using Prodigal v2.6.3 (option ‘-p’ for metagenome mode) (Hyatt et al., 2010). Third, the contigs analyzed were filtered to have a length of 20 kb or less. This criterion captured the twilight zone (5 kb to 11.6 kb) below the smallest isolated-tailed phage containing a major capsid protein, *Salmonella* phage ashtrithr (Luque et al., 2020; Olsen et al., 2020), and a control region above it (11.6 kb and 20 kb). The twilight zone was divided into micro genomes (< 5 kb), which had not been previously reported among HK97-fold entities, and mini genomes (5 to 11.6 kb).

The capsid phenotype was predicted using the genome-to-capsid model for HK97-fold protein shells (Lee et al., 2022; Luque et al., 2020). This included T=1 canonical capsids (hexagonal lattice) and extended capsids containing minor proteins, organizing the capsid proteins following the trihexagonal, snub hexagonal, and rhombihexagonal lattices (Twarock & Luque, 2019). In the absence of minor proteins, these non-canonical icosahedral capsids should be interpreted instead as elongated T=1 capsids (Luque et al., 2010; Luque & Reguera, 2010) or shrunk T=3 capsids (Bárdy et al., 2020).

### Genome annotation

The open reading frames (Data File S5) were annotated in three complementary steps (Figure 2A). First, high-level functional categories were predicted using the viral protein families protein language model, VPF-PLM (Flamholz et al., 2024). The VPF-PLM had been trained on the prokaryotic virus remote homologous groups (PHROGs) database (Terzian et al., 2021). Functions were assigned when obtaining likelihood scores greater than 60%; the highest-ranked function was selected. The method achieves a weighted F1-score (harmonic mean of the precision and recall) of 85% at this threshold for assigning correct high-level functional categories to viral proteins validated on ocean virome sequences (see annotations in Data File S6).

**Figure 2.**
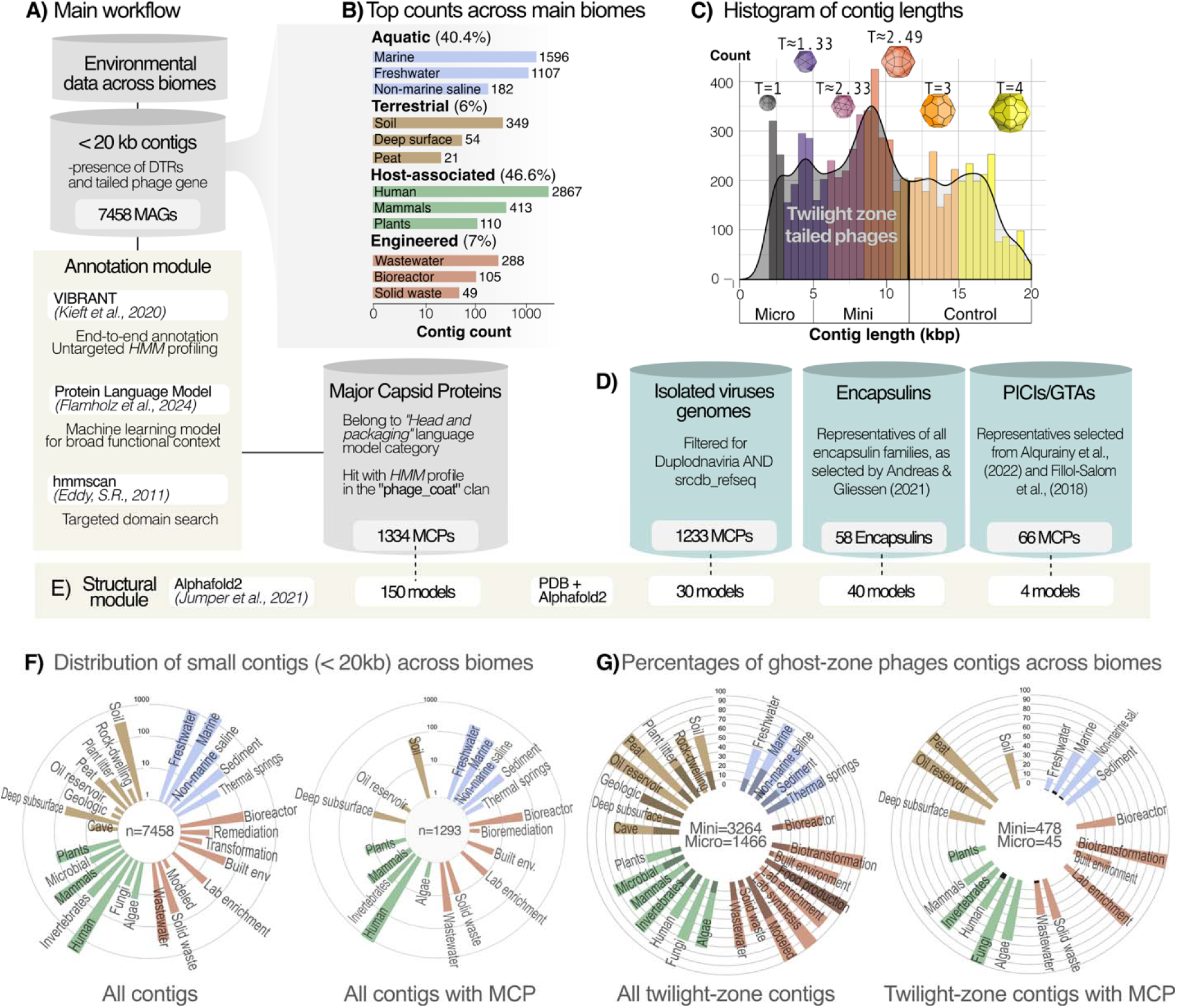
Discovery and comparison pipeline for twilight zone viruses. A) Flowchart illustrating the process from data acquisition to structural prediction. It is divided into three steps: filtering conditions, annotation, an major capsid protein identification. B) The bar plot displays the number of contigs (log scale) contributing from the top three biomes category (log scale) in each Level 1 category: Aquatic (blue), terrestrial (brown), host-associated (green), and engineered (red). C) Histogram of contig lengths. The solid line is a kernel density distribution with a bandwidth of 700. The colors are associated with different icosahedral capsids with the structures and T-numbers displayed on top. The vertical line represents the genome length threshold associated with the smallest isolated tailed phage genome, 11.6 kb, and the region below this value (the twilight zone) is shaded and includes mini (5 kb to 11.6 kb) and micro (< 5 kb) contigs. D) Overview of external datasets used for comparative analyses containing major capsid proteins (MCPs). E) Structural module indicating the number of structures predicted with AlphaFold2 or obtained from the Protein Data Bank (PDB). F) Circular bar plot representing the distribution of viral contigs across biomes, including all filtered contigs (left) and those with annotated MCP (right). The center includes the total number of contigs displayed (n). G) Similar plot to F but focuses on the Twilight Zone region. The center displays the number of mini and micro contigs.

Second, the VIBRANT (Virus Identification By iteRative aNnoTation) tool, which combines neural networks and hidden Markov models (HMMs) (Kieft et al., 2020), was applied to predict specific functions for each open reading frame. Version 1.2.1 of VIBRANT was executed with default parameters, which included curated HMM profiles from three major databases: KEGG (March 2019 release), Pfam (v32), and Virus Orthologous Groups (VOG; release 94) (see annotations in Data File S7).

Third, all predicted open reading frames, regardless of prior functional assignments, were scanned with hmmscan against the Pfam database (Mistry et al., 2021), clan CL0373 (“Phage-coat”), which contains curated HHM profiles for domains characteristic of HK97-fold. The clan comprised 17 HMM profiles, including viral capsid protein domains (14) and encapsulin shell domains (3). An additional viral capsid protein HMM profile (41798) from the efam database (Zayed et al., 2021) was included, as it represents a recently proposed HK97-fold MCP lineage identified using protein language models (Flamholz et al., 2024). Hits were accepted at a cutoff of E-value < 10 ³. The HMM profile list and annotations are provided in Data File S8.

After domain-based identification of HK97-fold proteins, a set of 60 viral genomes (encompassing 639 open reading frames) was analyzed in detail for quality control when Pfam CL0373 (“Phage-coat”) domains were detected but not classified as MCPs by either VIBRANT or VPF-PLM. It also included genomes with a high proportion of unannotated proteins. These were manually annotated using the HMM-based remote protein homology detection server HHpred (Söding, 2005). Queried annotation libraries included Pfam-A (v37), NCBI Conserved Domains (v3.18), and PHROGs (v4) with default server parameters and a MAC realignment threshold of 0.3. The Protein Data Bank (PDB) library was included to enhance structural evolutionary insights. Hits below 20% probability were discarded. For each open reading frame, the top hit was selected unless shorter than a subsequent hit of comparable significance. Functions were labeled according to statistical confidence: very high (>99%, ***), high (95– 99%, **), moderate (70–95%, *), and low (50–70%, ?). Annotations below 50% probability were considered hypothetical. Manual annotations took precedence when conflicting with automated assignments. The HHpred annotations are provided in Data File S9.

### Phylogeny of environmental HK97-fold major capsid proteins

The amino-acid sequences of the environmental HK97-fold proteins (Data File S10) were clustered using a sequence-based phylogenetic inference of the HK97-fold major capsid proteins.

The multiple sequence alignment was performed using MAFFT v6.240 (Katoh, 2002). The alignment of conserved cores was optimized by the L-INS-i algorithm, which adds a generalized gap cost to accommodate variable regions (Katoh & Toh, 2008). The alignment was refined, and artifacts were reduced by increasing the number of iterative cycles up to 1000 using the--maxiterate parameter. Gap penalties were adjusted with gap thresholds of 0.05 and 0.15, which are standard when aligning highly variable multi-domain viral proteins (Kazlauskas et al., 2019). Phylogenetic inference was performed using maximum likelihood as implemented in IQ-TREE 2 v2.3.6 (Minh et al., 2020). The best-fit substitution model was based on the Bayesian Information Criterion (BIC) and obtained with ModelFinder, integrated into IQ-TREE 2. The tree was inferred with ultrafast bootstrap approximation using 1000 replicates, and the default consensus tree was generated. The outputs, including the maximum-likelihood tree, log file, and likelihood distances, are provided in Data File S11. The tree was visualized and annotated using iTOL v6 (Letunic & Bork, 2024), overlaying the biome of origin and genome length. The groups derived from the phylogeny were explored by partitioning the tree into clusters using TreeCluster v1.0.3 (Balaban et al., 2019), which generates biologically meaningful clusters based on tree-based distances. Clustering was performed using the avg_clade method, which groups leaves into clusters so that the average pairwise distance between leaves does not exceed a specified threshold while ensuring each cluster forms a distinct clade. A threshold of 0.001 was applied to identify highly specific groupings.

### Comparative analysis across HK97-fold proteins

*Reference HK97-fold protein database.* The HK97-fold proteins obtained from environmental samples were compared with a reference database composed of HK97-fold proteins from curated sources, including viral genomes from NCBI RefSeq, viral-related mobile elements (gene transfer agents, GTAs, and phage-inducible chromosomal islands, PICIs), and encapsulins (Data Files S12 and S13). Viral MCP sequences belonging to the *Duplodnaviria* group were retrieved from the NCBI Viral Genomes Resource (Brister et al., 2015) in December 2023 using the Boolean query (“MCP” OR “Major Capsid Protein”) AND “viral” NOT “human” AND “refseq”, filtering to select only those belonging to complete genomes. These sequences were further confirmed to contain HK97-fold domains using hmmscan against the Pfam clan CL0373 (“Phage-coat”) with an E-value cutoff < 10 ³. The resulting validated sequences constituted the final reference dataset. The MCPs encoded by PICIs were obtained using the NCBI identifiers reported in PICIs-related publications (Alqurainy et al., 2023; Fillol-Salom et al., 2018; Penadés & Christie, 2015), yielding 66 MCPs. The set of encapsulin sequences was obtained from a recent study that expanded the encapsulin universe (Andreas & Giessen, 2021), yielding 58 proteins from four encapsulin families: family I (51 proteins), family II (30), family III (7), and family IV (13).

*Estimating packaging capacity of encapsulins.* To compare the packaging capacities of viral entities and encapsulins, the internal volume of encapsulins forming T=1 (PDBs: 3DKT, 6X8T), T= (4PT2, 2E0Z), and T=4 (6NJ8) protein shells was measured using the Blob and Surface tools in ChimeraX (Meng et al., 2023). The encapsulin structure was built from the asymmetric unit by applying icosahedral symmetry operations, a density map was generated, and the contour level was adjusted to define the capsid boundaries. The Blob Picker tool selected the inner capsid surface and measured the enclosed volume (Figure S1). The encapsulin packaging capacity was projected using the average density of double-stranded DNA in tailed phage capsids, ~0.5 kbp/nm^3^ (Luque et al., 2020; Podgorski et al., 2023; Stone et al., 2019).

*Pfam domain analysis*. The HK97-fold hidden Markov model (HMM) profiles captured in the phage_coat Pfam clan (CL0373) were mapped to the major capsid protein sequences in the environmental groups (micro, mini, and control) and the reference HK97-fold database. For each protein, the phage_coat coverage was calculated as the number of residues related to the non-overlapping Pfam domains identified divided by the protein’s amino-acid sequence length. The encapsulin enrichment of a protein was obtained as the fraction of the phage_coat coverage associated with a Pfam encapsulin domain. This domain analysis enabled the identification of viral MCPs exhibiting Pfam domains typically associated with encapsulins.

*Protein-domain co-occurrence analysis*. Proteins containing more than one phage_coat Pfam domain were used to build a co-occurrence network using the Python NetworkX library (Hagberg et al., 2008). Each node corresponded to a distinct Pfam domain. Edges were drawn between nodes (Pfam domains) co-occurring in the same protein. The edges were categorized as canonical when connecting viral-viral or encapsulin-encapsulin Pfam domains and non-canonical when connecting viral-encapsulin Pfam domains. The placement of the nodes was determined with a spring layout algorithm. Nodes with a larger degree of connectivity (number of connections) were positioned closer to each other. The algorithm adjusted the position of the nodes iteratively until equilibrium was reached, minimizing the overall spring tensions in the network. Node size was proportional to that node’s degree of connectivity, scaled between 100 and 500 units. The networks for RefSeq, control, mini, and micro subsets were compared.

*Structural phylogeny*. Viral phylogenetic analyses based on fold-centric structural comparisons have been used to elucidate deep evolutionary relationships among viruses and cells (Nasir & Caetano-Anollés, 2015). Due to the lack of protein sequence similarities, the evolutionary relationship of HK97-fold proteins was inferred based on their protein structures.

First, HK97-fold candidates were selected. This included 220 major capsid proteins (MCPs) from environmental contigs, ensuring representation across the length-based groups of twilight viral genomes (micro and mini) and control environmental viral contigs. This set included 113 MCPs enriched in encapsulin domains. An additional set of 147 MCPs was selected from the reference HK97-fold database. This included 16 proteins used in HK97-fold proteins whose structure had been obtained experimentally (and deposited in the Protein Data Bank, PDB), and had been used in prior evolutionary studies (Krupovic & Koonin, 2017), 14 MCPs from isolated viruses encoding encapsulin-enriched HK97-fold domains identified in this study, 6 encapsulin shell proteins from the structures available in the Protein Data Bank (PDB), encompassing families I, II, and IV, 4 MCPs from PICIs, and 1 MCP encoded in a gene transfer agents (GTA), which are taxonomically restricted to *Rhodobacter* (Bárdy et al., 2020). Second, for protein structures not available in the PDB, we predicted them using AlphaFold2 in ColabFold (Jumper et al., 2021). Structures were evaluated using the pLDDT (predicted Local Distance Difference Test) confidence metric. Folded protein structures scoring above 40 were retained for phylogenetic analysis. The empirical and predicted protein structures, including the pLDDT scores, are provided in Data File S14. Third, the structural phylogenetic tree for the HK97-fold proteins was constructed using FoldTree, which provides an alternative to sequence-based phylogenetic methods for distantly related proteins with a conserved structure (Moi et al., 2023; Mutti et al., 2025). FoldTree utilizes local structural alphabets that capture key features of protein structures, thereby avoiding issues associated with global structural metrics. Default parameters were used and relied on FoldSeek for the all-vs-all structural comparison (van Kempen et al., 2024). Structural tree files are available in Data File S15.

## RESULTS

### Twilight viral genomes are frequent, and a fraction contain identifiable HK97-fold proteins

There were 4,730 putative complete viral genomes in the twilight zone (< 11.6 kb) containing direct terminal repeats (DTRs) and predicted to be related to tailed-phage-like viruses (Figure 2C), representing 63% of the total viral contigs when including the control region (11.6 – 20 kb), which contained 2,728 viral contigs. The twilight viral genomes were frequent across biomes (Figure 2F-G). However, the fraction of identified HK97-fold major capsid proteins (MCPs) was significantly smaller in the twilight zone (11% or 536 MCPs) than in the control region (29% or 798). Most twilight viral genomes containing MCPs were from the human gut (46% or 245). A few biomes yielded more MCPs from twilight viral genomes than in the control zone (Table S1). This included non-marine saline, invertebrates, peat, and oil reservoir biomes (Figure 2G).

### Significant encapsulin domain enrichment found in MCPs from mini viral genomes

The working hypothesis in the study was that HK97-fold capsid proteins from twilight viral genomes will be more closely related to encapsulin proteins. This was tested by comparing the presence of encapsulin domains in viral HK97-fold proteins (Figure 3). The analysis mapped the HK97-fold’s hidden Markov model (HMM) profiles from the phage_coat Pfam clan to assess the protein’s phage_coat coverage and the fraction of coverage associated with encapsulin domains (Figure 3A). The phage_coat amino-acid coverage decreased for MCPs from smaller viral genomes as expected (Figure 3B). The MCPs in micro genomes (< 5 kb) displayed the lowest coverage, with a median of 50% of the MCP sequence. The mini and control genomes displayed a median coverage of 70%. The RefSeq group displayed the most extensive coverage, with a median of 95%, as expected. The viral genomes displaying the most substantial enrichment in encapsulin domains were from twilight genomes as hypothesized (Figure 3C). However, the hits concentrated, unexpectedly, in MCPs from mini genomes (5 kb to 11.6 kb) instead of micro genomes (< 5 kb), where none of the 45 identified MCPs were encapsulin-enriched. Nearly 19% of MCPs in mini genomes (93 out of 491) contained encapsulin domains. Among those, more than half (57 out of 93) displayed encapsulin domains as their only phage_coat profile, while the remaining 36 displayed fractions between 0.14 and 0.55. The encapsulin-domain enrichment of MCPs from mini genomes was significantly higher than in the environmental control group (11.6 kb to 20 kb), where only 2.5% (20 out of 798) of the MCPs contained encapsulin domains, although 4 displayed an encapsulin domain fraction of 1 (Figure 3C). In the RefSeq group, only 1.9% (17 out of 910) MCPs contained a trace of encapsulins domains. All their enriched MCPs displayed encapsulin fractions below 0.5, and were found in genomes of lengths around ~50 kb and ~100 kb (Figure 3D and Data File S16), except for one MCP with fraction 1 from the deep-sea archaea virus Verda V2 (NC_074645.1), a metagenomically assembled genome (19.5 kb) in the fringe zone (Medvedeva et al., 2022). The Refseq genomes encoded hallmark HK97-like MCP, portal, and tail proteins. No HK97-fold proteins from the viral mobile elements (GTAs or PICIs) displayed encapsulin domains.

**Figure 3.**
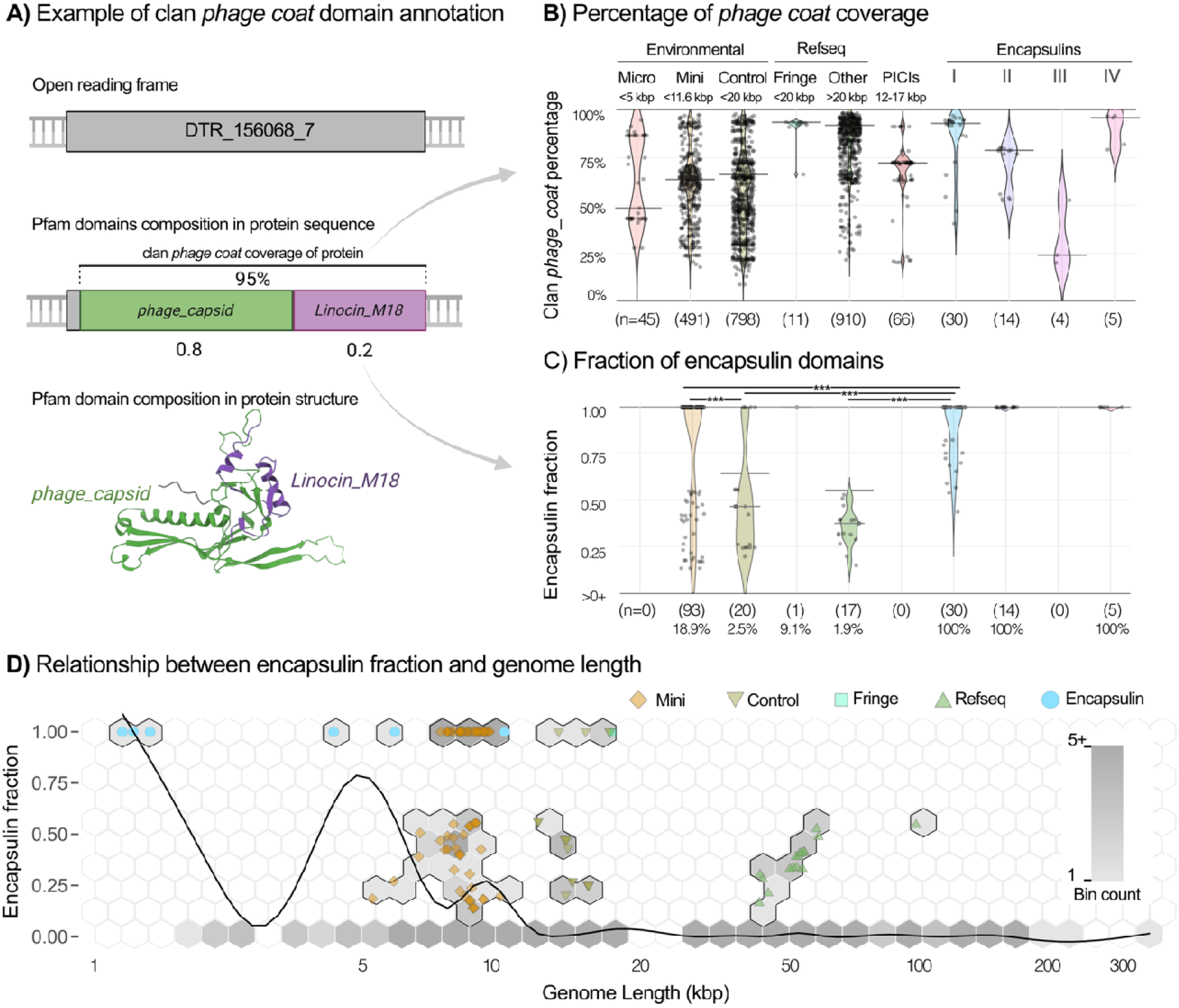
Encapsulin domain analysis across viral entities. A) Schematic representation of the major capsi protein domain analysis. The open reading of an annotated major capsid protein is displayed in grey. The protei domains identified are displayed in different colors, indicating the names of the specific family domains. The blac horizontal bar covers the residues captured by the combined phage_coat clan domains, and the percentage of coverage is displayed. The numbers under the domains indicate the fraction of each domain. The predicted 3D protein structure is colored based on the sequence associated with each phage_coat domain identified. B) Percentage of phage_coat coverage plotted for the MCPs for viral and encapsulin groups. The black dots are the values for each representative. The horizontal bars represent the median. Each violin plot is displayed in a different color based on the group, with names on top. The number of members (n) in each group is displayed in parentheses at the bottom. C) Fraction of the encapsulin domains observed in the major capsid proteins for each group, following the same logic as in panel B). The symbol >0% in the y-axis indicates that only MCPs with an encapsulin domain fraction above zero are displayed. The horizontal bars indicate the groups that were compared statistically, displaying stars if they were different statistically. D) Scatterplot of encapsulin fraction versus genome length for those elements with an encapsulin signal (>0%). Each group is represented with a different symbol (see legend). The colors correspond to the same schema as in B) and C). For the encapsulins, the projected genome length corresponds to the calculated packaging capacity. The hexagonal net is a hexbin plot showing the density of data points on a grey scale, as indicated by the color bar. It includes the members that contain no encapsulin domain signal.

The prevalence of encapsulin domains across viral MCPs decreased with the viral genome length (Figure 3D). The encapsulin-enriched twilight mini genomes (5 kb - 11.6 kb) occupied the genome length region between the two largest encapsulin systems, which displayed a projected approximated genome capacity of 5 kb and 10 kb, respectively. The few encapsulin-enriched MCPs in the control group were associated with genome lengths ranging from 13 kb to 19 kb, which is above the maximum projected packaging capacity of known encapsulins. The even smaller fraction of viral genomes from RefSeq containing MCPs with encapsulin domains displayed genome lengths ranging from 19 kb to 58 kb, except for one virus with a 103 kb genome length (Figure 3D). The list of identifiers corresponding to contigs with encapsulin enriched MCPs is included in the supplementary material Data File S16.

### A consistent clade of encapsulin-enriched viral genomes is temperate and tailless

The phylogeny of major capsid proteins (MCPs) revealed that encapsulin-enriched MCPs formed 10 clades and 10 singletons (Figure 4A). One of the clades was dominant, containing 57% (75) of the encapsulin-enriched MCPs. Their functional analysis of the viral genomes revealed common characteristics (Figures 4B). They originated from host-associated (human gut) samples. Their average genome length was 8.9 (± 0.6) kb. Genes of unknown function accounted on average for 24% (16% - 32 %) of the genome or 1.7 kb (1.0 - 2.4 kb). Structural genes covered an average of 22% (16% - 28%) of the genome or 1.5 kb (1.0 kb - 2.0 kb) (Figure 4C). The structural proteins were MCP and scaffold. Transcription regulation genes covered an average of 21% (15% - 27%) or 1.4 kb (1.0 kb - 1.8 kb). They included RNA polymerase sigma factors, transcriptional repressors, and often quorum-sensing proteins. DNA and RNA metabolism genes comprised 17% (10% - 24%) of the contig length or 1.2 kb (0.8 kb - 1.6 kb), including RepA, DNA helicase, or primosomal DNA I. Lysogenic genes represented 16% (12% - 20%) of the genome or 1.1 kb (0.9 kb - 1.3 kb), including integrase and, in some cases, excisionase. A minor fraction of the genome, 0.4% (0% - 2.4 %) or 0.026 kb (0 kb - 0.126 kb), encoded auxiliary metabolic genes, such as ribosomal proteins, membrane receptors, or formin-like proteins. No terminases or packaging genes were found. No portal, tail connector, or structural tail genes were detected. The only trace of a tail-related function among encapsulin-enriched genomes corresponded to a single open reading frame (DTR_497289_3), annotated as a tail tape measure protein.

**Figure 4.**
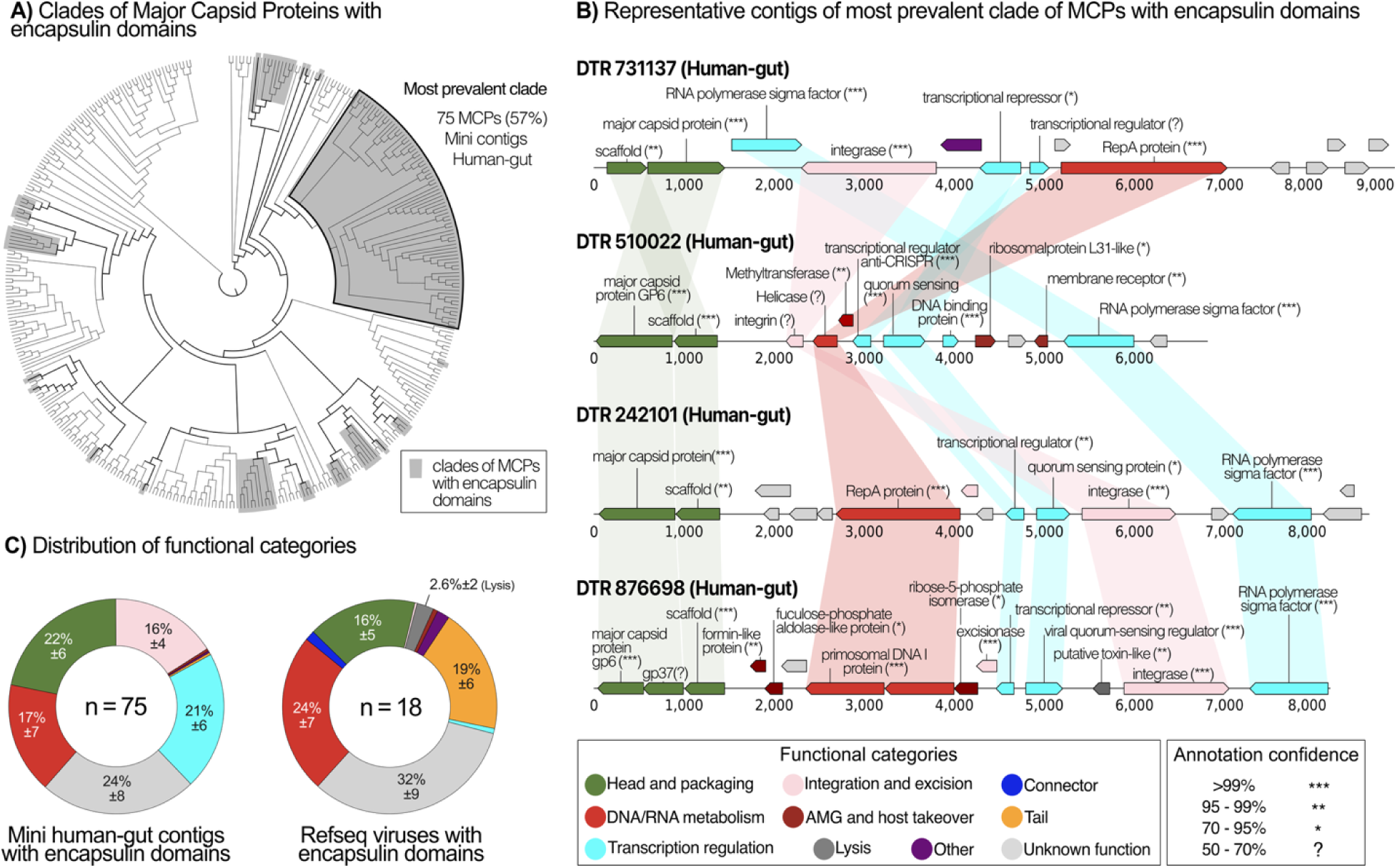
Genome composition of encapsulin-enriched mini viral entities. A) Gene maps of representative genomes from the encapsulin-enriched mini virus group. The horizontal line represents the putative genome, and the numbers are the length in bp. The open reading frames are displayed as thick-colored arrows. The colors correspond to high-level genome functions listed in the legend on the top right of the figure. The genes annotated successfully include the predicted function as a label with its statistical significance in parentheses (see methods). Each genome also includes the identifier (DTR), which all come from human-gut samples. B) Distribution of functional categories for the encapsulin-enriched mini viral group (left) and the encapsulin-enriched representatives from RefSe (right)among viruses with encapsulin fractions. The colors are related to the functional categories, as in panel A). The sections of the pies include the average percentages.

The RefSeq phages containing encapsulin domains in their MCPs were also temperate (containing lysogenic genes). They primarily infect bacteria in the microbiome of mammals, including humans, and a few infect archaea from the order Poseidonales. Their genome composition encoded tail genes as expected, comprising an average of 19(± 6)% of the genome (Figure 4C and Data File S17). The twilight viral genomes that did not encode encapsulin-enriched MCPs did not form a homogenous group. However, tail genes were also uncommon (Table S2). Only 4.3% (23 out of 536) of twilight genomes with MCPs encoded a tail gene, and the majority (20 out of 23) encoded just one tail gene. Among the control group, 12.2% (97 out of 798) encoded tail genes, but the control genomes containing encapsulin-enriched MCPs did not encode tail genes.

### Bidirectional evolution of HK97-fold viruses and encapsulins

The evolutionary analysis of the HK97-fold proteins indicated bidirectional evolution of HK97-fold viruses and encapsulins. This conclusion was reached by combining two complementary analyses described below, one based on the co-occurrence of viral and encapsulin HK97-fold Pfam domains in proteins and the other on the large-scale structural phylogenetics of HK97-fold proteins.

Regarding the co-occurrence domain analysis, proteins were categorized as viral or encapsulin based on their genomic origin. A sub-category was added based on the provenance of the HK97-fold HHM profiles from the phage_coat Pfam clan, including viral, encapsulin, or mixed, when viral and encapsulin domains were found in the same protein. These yielded representatives for five out of the six possible groups (Figure 5A). Viral HK97-fold MCPs (2,366) formed one dominant group with only viral domains (n = 2,235 or 94%) and two minor groups: one combining viral and encapsulin domains (n = 69 or 3%) and the other containing only encapsulin domains (n = 62 or 3%). HK97-fold encapsulins (55) formed two groups: the dominant group displayed only encapsulin domains (n = 38 or 69%), and the smaller one combined viral and encapsulin domains (n = 17 or 31%). No HK97-fold encapsulins were found to contain only viral domains. The co-occurrence analysis of phage_coat domains revealed that the MCPs from small uncultured genomes (control and mini) were necessary to fully connect the network of HK97-fold protein domains (Figures 5B and S2). They also significantly increased the robustness of the existing non-canonical viral-encapsulin connections present in the RefSeq subset (Figure S2). The Linocin_M18 and Srpl-like encapsulin domains exhibited extensive non-canonical connections with viral domains, displaying 88 and 8 connections, respectively (Figure 5B). Linocin_M18 (Family I encapsulins) was primarily connected with phage_capsid domains (35 out of 88), while Srpl-like (Family II encapsulins) connect mostly with Gp7 domains (6 out of 8).

**Figure 5.**
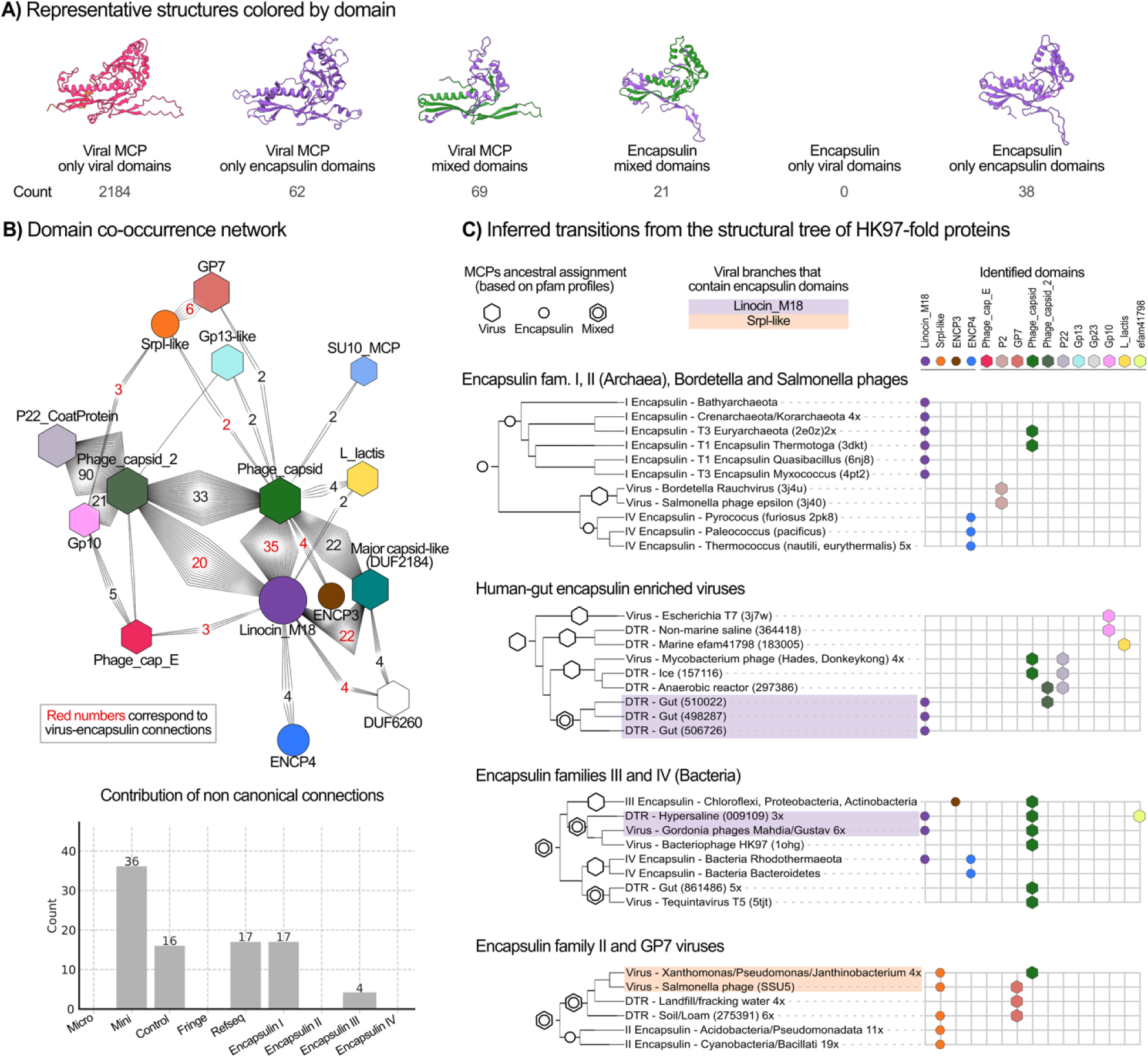
Structural phylogenetics and domain co-occurrence network. A) Representative structures colored b Pfam domain. Six structural groups are shown, and the number of hits in each group is provided in the row labele as Counts. Each model is colored according to its Pfam profiles (see legend in C); mixed-domain proteins displa multiple colors. B) Pfam domain co-occurrence network. Nodes represent individual Pfam profiles (color-coded as in C); node size scales with total degree. Edges link profiles found in the same protein, with edge labels showing co-occurrence counts (red numbers denote counts of non-canonical virus–encapsulin links). Below, a bar plot breaks down the categories contributing to non-canonical connections (micro, mini, control, fringe, RefSeq, Encapsulins I– IV), highlighting the predominance of the “Mini” group. C) Inferred domain transitions on the rooted structural phylogeny. Shown are only branches containing both virus-and encapsulin-associated profiles. Tip symbols mark Pfam domains presence (hexagon = virus, circle = encapsulin; combined shapes = mixed-domain proteins). Symbols placed at nodes or mid-branches directly on top of the tree indicate inferred ancestral assignments based on domain prevalence. Shaded boxes highlight viral clades enriched in encapsulin domains (Linocin_M18, SrpI-like). To the right of each clade, colored symbols denote the specific Pfam domains present.

The structural phylogenetic analysis revealed different instances where viral MCPs displayed a close evolutionary relationship with encapsulin proteins. These instances were present in three clades, I, II, and VII (Figure 5C), out of the eight clades identified in the structural evolutionary analysis of HK97-fold MCPs (Figure S3). Clade I grouped HK97-fold proteins from encapsulin Family II, terrestrial (soil/fracking) viral genomes from the control group, and isolated phages infecting predominantly soil bacteria (Xanthomonas and Janthinobacterium) and host-associated bacteria. Srpl-like encapsulin domain and GP7 virus domains were widespread in this clade. Encapsulins are deeper in the tree, and Srpl-like domains are the most dominant in the Clade; thus, an encapsulin ancestor is more likely. Clade II contained two instances of virus-encapsulin evolution. One involved bacterial encapsulins from Family IV (bacteriodetes-like groups), mini DTRs from the gut, and coliphage T5. It is unclear if the ancestor in this group is encapsulin or viral. The other group involved encapsulin from Family III, characteristic of soil bacteria, mini DTRs from hypersaline environments, and isolated phages from soil bacteria. The presence of Phage_capsid viral domains indicates that encapsulin from Family III might have derived from a viral HK97-fold ancestor. Linocin_M18 encapsulin domains were also common but less frequent as Phage_capsid domains in Clade II. The fourth instance was observed in Clade VII, which grouped archaeal encapsulins from Family IV, found in thermophiles, and MCPs from phage infecting *Salmonella* and *Bordetella spp.*, which are hosts that can replicate intracellularly in warm-blooded animals like humans. The proximity of Clade VIII, which only contains encapsulin, suggests a potential encapsulin-like ancestor for the viruses in Clade VII.

## DISCUSSION

### Procapsid as the evolutionary nexus and ancestor of HK97-fold compartments

The structural phylogenetic analysis of HK97-fold proteins revealed multiple instances where either viruses derived from encapsulins or vice versa, summarized in Figure 6A. This included at least one encapsulin group that likely derived from HK97-fold viral MCPs (Family III in Clade II), one virus group that likely derived from encapsulins (Clade VII), and two instances with mixed virus-encapsulin derivations (Clades I and II) (Figures 5C and S3). Thus, modern HK97-type viruses and encapsulins seem to have evolved from each other, bidirectionally, multiple times, contrary to the current paradigm (Andreas & Giessen, 2021; Krupovic et al., 2019). This bidirectional evolutionary paradigm is further supported by the fact that HK97-fold protein domains, traditionally assigned to viruses and encapsulins, form a fully connected network, as revealed by the addition of HK97-fold proteins from small uncultured viruses (Figures 5A, 5B, and S2).

**Figure 6.**
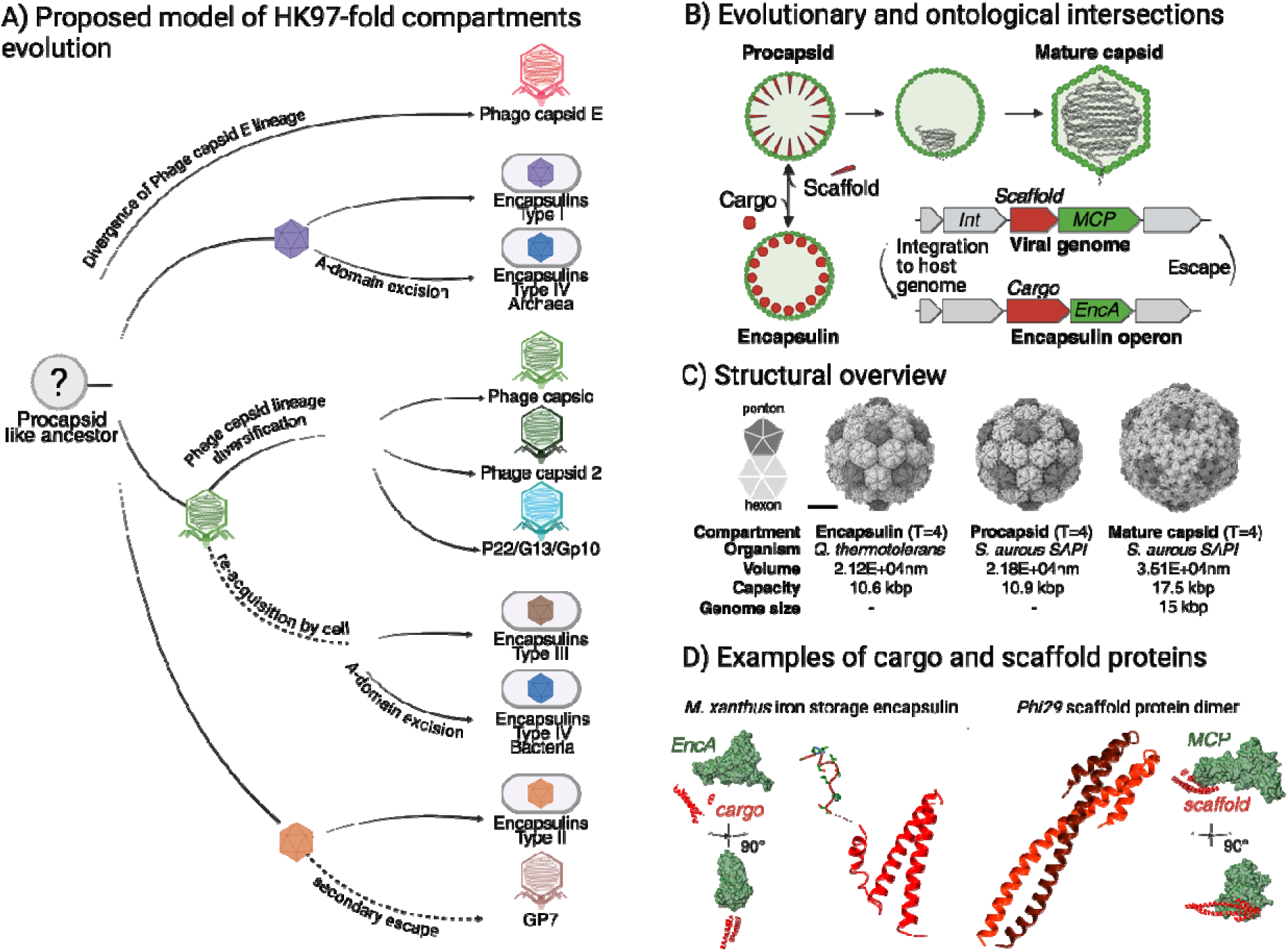
Model of HK97 fold protein evolution. A) HK97 fold evolutionary model enhanced with twilight-zone phages. This model posits multiple transitions between viral MCPs and encapsulin, suggesting that encapsulins di not emerge once as a monophyletic group of proteins, but rather through independent transitions from viral ancestors, and vice versa. A procapsid-like ancestor to viral and cellular HK97-fold compartments was proposed t link the biochemical and genetic delivery roles of the known diversity. B) Evolutionary and ontological intersections between viral MCPs and encapsulins. This panel illustrates the parallel between scaffolding proteins in viral capsids and cargo proteins in encapsulins, implying that both play a functionally equivalent role in determining the final assembly and purpose of the compartment. The procapsid, an immature and empty stage of viral capsids, coul represent a bifurcation point in HK97-fold evolution, where modifications to its function led to the divergence of viral capsids and encapsulins. Showed structures correspond to PDB IDs: 7S8T (Ferritin-like encapsulin cargo system) and 8F2M (Phi-29 MCP bound to scaffold protein dimer). C) Structural comparison of HK97-fol compartments. The structural resemblance between encapsulins (PDB: 6NJ8) and viral procapsids (PDB: 7RWZ) is highlighted. Additionally, the comparison emphasizes a key constraint in viral evolution: at the procapsid stage, the internal volume is insufficient to accommodate the genome that the mature capsid will later encapsulate, necessitating further structural rearrangements. D) Representative structures of an encapsulin cargo protein from Myxococcus xanthus (EncA; PDB 7S8T) and the bacteriophage Φ29 scaffold protein dimer (PDB 8F2M), each adopting a “hinge-like” fold.

Bidirectional evolution has been described in viruses, for example, in how helicases in viral lineages that form capsids with the jelly-roll fold have transitioned between plasmids and viruses (Kazlauskas et al., 2019). The question is what molecular mechanism would explain the recurrent bidirectional evolution inferred here for encapsulins and viruses. One compelling scenario is that viral procapsids have facilitated such transitions (Figure 6B). This argument is based on the structural similarity of encapsulins and procapsids (Figure 6C), the co-assembly and structural parallels of cargo and viral scaffold proteins (Figure 6D), and the presence of lysogenic genes in the well-defined mini viruses encoding encapsulin-enriched HK97-fold protein discovered in this study (Figures 3C and 4).

Encapsulins and procapsids are structurally more similar to each other than procapsids are to their mature state (Figure 6C). This comparison was possible thanks to the existence of a few high-resolution T=4 architectures found among encapsulins as well as HK97-fold procapsids and mature capsids (Figure 6C). This includes the unrelated encapsulin from *Bacillus thermotolerans* (Giessen et al., 2019) and the *Staphylococcus aureus* Pathogenicity Island 1 (SaPI1), which repurposes the HK97-fold protein from phage 80α (Dearborn et al., 2017). The encapsulin and procapsid structures enclosed comparable volumes around ~2.15·10 nm³ with an equivalent packaging capacity of ~10.75 kbp, which was 60% smaller than the internal volume (~3.51·10 nm³) and packaging capacity (~15.2 kb) of the mature capsid (Figure 6C). The expansion in volume from the procapsid to mature capsid stage is common across *Duplodnaviria* phages (Figure S4) (Hua et al., 2017; Veesler & Johnson, 2012). Encapsulins and procapsids form shells that are more porous and display bulgier capsomers (pentamers and hexamers) than mature capsids (Giessen & Silver, 2017; Hryc et al., 2017; Selivanovitch et al., 2021; Wu et al., 2016). The pores enable encapsulins to transport reactants (ions or other molecules) necessary for biochemical functions (Chmelyuk et al., 2023; Ross et al., 2022; Williams et al., 2018). The pores in procapsids have facilitated their use as biochemical reactors in engineering applications (Selivanovitch et al., 2021; Wang & Douglas, 2022). A second important observation between encapsulins and procapsids is that they both co-assemble with proteins, respectively, cargo and scaffold, which determine their function (Figure 6B)(Giessen, 2016; Sutter et al., 2008). Both cargo and scaffold form proteins that are enriched in alpha-helices and often include disordered regions that are difficult to characterize empirically (Huet et al., 2023). Their structural comparison indicates clear parallels, in particular, the presence of a helix-loop-helix motif (Figure 6D).

The third argument in support of the bidirectional evolutionary molecular transition of encapsulins and viruses via procapsids is based on the well-defined group of mini viral genomes described here, which contained the most encapsulin-enriched HK97-fold proteins (Figures 3C and 4). This group is characterized by the presence of lysogenic genes, indicating the potential for genomic integration in the host genome (Howard-Varona et al., 2017; Luque & Silveira, 2020). The integration in the host would increase the likelihood of viral procapsids incorporating cargo-like protein to become encapsulins. It could also facilitate encapsulins to recombine with scaffold proteins, leading to the emergence of encapsulin-derived viruses (Figure 6A and 6B). Conversely, an encapsulin shell might reacquire genome packaging function through recombination, inserting a small scaffolding protein. This change would repurpose a metabolic compartment back into a viral-like entity. Together with the hinge like fold shared by scaffold and cargo (Figure 6D), these scenarios illustrate a flexible, bidirectional model in which the identity of the internal “hinge” protein determines the shell’s role. The fact that the few RefSeq phages containing encapsulin-enriched HK97-fold domains are lysogenic further supports this scenario (Figure 3). The larger capsid size of these phages raises the possibility that related encapsulins may form larger architectures, T > 4, which are not among the canonical encapsulins, T ≤ 4 (Chmelyuk et al., 2023; Giessen & Silver, 2017).

This virus-encapsulin transition is consistent with frameworks considering that viruses function as integral components of larger biological systems (holobionts), facilitating extensive genetic and functional exchanges with cellular hosts (Caetano-Anollés, 2024). Given the similarities between encapsulins and procapsids, if viral genomes or the hosts encode cargo-related proteins that are co-assembled in procapsids, then procapsids could carry biochemical functions during viral infections. Given the parallels between viral scaffold proteins and encapsulin cargo proteins, it is not impossible that scaffold proteins could retain biochemical functions. The fact that viral infections often generate empty capsids or procapsids (Laurenceau et al., 2020; Subramanian et al., 2025) makes the potential biochemical role of procapsids a subject for scrutiny moving forward, which would require a complete revision of the role of viral capsids during infections.

### Lack of tails and portal in HK97-fold twilight viruses

The well-defined group of encapsulin-enriched mini viral genomes lacked portal proteins and tail genes (Figure 3). But encoded replicase RepA-like genes are commonly found in the replication of ssDNA viruses and plasmids (de la Higuera et al., 2020; Kazlauskas et al., 2019), including the plasmid phase of HK97-type phages *P4* and *N15* (Ravin, 2015). In ssDNA viruses, the activity of RepA is either directly or indirectly associated with the packaging of the genome in the empty capsid through 3-fold or 5-fold pores (Mankertz & Hillenbrand, 2001; Sonntag et al., 2011). The discovery of ssDNA viruses with small genomes (< 11 kb) in a structural viral lineage (double-jelly fold) traditionally associated with dsDNA viruses (Kejzar et al., 2022; Krupovic et al., 2024; Laanto et al., 2017; San Martín & Van Raaij, 2018) makes the existence of portal-less and tail-less HK97-fold ssDNA viruses something possible. This clade could represent a case of HK97-fold viruses packaging ssDNA genomes, departing from one of the canonical traits of *Duplodnaviria* viruses (Koonin et al., 2020).

Among isolated HK97-fold viruses, satellite lifestyles seem more common among those with relatively short genomes (12 kb to 20 kb) and HK97-type mobile elements (11 kb to 18 kb) (Agarwal et al., 1990; deCarvalho et al., 2023; Penadés et al., 2025; Pourcel et al., 2024). The need for helper viral elements or unusual host conditions would explain the difficulty in isolating the putative twilight HK97-fold viruses discovered in this study. The presence of tail genes in a small fraction of twilight viruses may be a remnant of a more canonical tailed phage structure and replication, possibly in combination with helper viruses. Nonetheless, it is important to highlight that the characteristic small ssDNA viruses (< 6 kb) in the *Microviridae* family, which form tailless viruses, can generate a dynamic tail when infecting (Kirchberger & Ochman, 2023; Sun et al., 2013, 2014). Therefore, the few tail genes observed in the HK97-fold twilight viruses could operate similarly to dynamic tails instead of the canonical *Duplodnaviria* tails.

## CONCLUSIONS

The molecular relationship revealed between HK97-fold proteins from small viral genomes and encapsulins supports a change of paradigm in the evolution of HK97-fold compartments, favoring a bidirectional evolution where the procapsid represents the most likely nexus and common ancestor. The potential genomic and biochemical functions of procapsids make this system particularly interesting for the evolution of early life. It also opens up the possibility that viral procapsids may have the capability to carry metabolic functions, which would necessitate a reevaluation of viruses in the tree of life (Moreira & López-García, 2009). The discovery of a well-defined clade of encapsulin-enriched HK97-fold twilight viruses opens the doors to follow-up research to confirm further the findings and predictions described here.

## ACKNOWLEDGMENTS

The authors acknowledge Sean Benler, Cynthia Silveira, Kevin Collins, and Neil Rosser for fruitful discussions throughout different stages of the project. A.A. and A.L. acknowledge the support from the Gordon and Betty Moore Foundation award GBMF9871 (https://doi.org/10.37807/GBMF9871) and the National Science Foundation award #2424579. The work conducted by the U.S. Department of Energy Joint Genome Institute (https://ror.org/04xm1d337), a DOE Office of Science User Facility, is supported by the Office of Science of the U.S. Department of Energy, operated under Contract No. DE-AC02-05CH11231.

## Notes

https://drive.google.com/drive/folders/1s4JYU9VXwNfHClpkb8KhwEveDSAKeLWu?usp=drive_link

### Competing Interest Statement

The authors have declared no competing interest.

### Summary of Updates

Corrected the order in which authors appear in the bioRxiv system.

## REFERENCES

Agarwal, M., Arthur, M., Arbeit, R. D., & Goldstein, R. (1990). Regulation of icosahedral virion capsid size by the in vivo activity of a cloned gene product. Proceedings of the National Academy of Sciences, 87(7), 2428–2432. 10.1073/PNAS.87.7.2428

Aksyuk, A. A., Bowman, V. D., Kaufmann, B., Fields, C., Klose, T., Holdaway, H. A., Fischetti, V. A., & Rossmann, M. G. (2012). Structural investigations of a *Podoviridae streptococcus* phage C1, implications for the mechanism of viral entry. Proceedings of the National Academy of Sciences, 109(35), 14001–14006. 10.1073/pnas.1207730109

Alqurainy, N., Miguel-Romero, L., Moura de Sousa, J., Chen, J., Rocha, E. P. C., Fillol-Salom, A., & Penadés, J. R. (2023). A widespread family of phage-inducible chromosomal islands only steals bacteriophage tails to spread in nature. Cell Host & Microbe, 31(1), 69–82.e5. 10.1016/j.chom.2022.12.001

Andreas, M. P., & Giessen, T. W. (2021). Large-scale computational discovery and analysis of virus-derived microbial nanocompartments. Nature Communications, 12(1), 4748. 10.1038/s41467-021-25071-y

Balaban, M., Moshiri, N., Mai, U., Jia, X., & Mirarab, S. (2019). TreeCluster: Clustering biological sequences using phylogenetic trees. PLOS ONE, 14(8), e0221068. 10.1371/journal.pone.0221068

Bárdy, P., Füzik, T., Hrebík, D., Pantůček, R., Thomas Beatty, J., & Plevka, P. (2020). Structure and mechanism of DNA delivery of a gene transfer agent. Nature Communications, 11(1), 3034. 10.1038/s41467-020-16669-9

Becerra, A., Delaye, L., Islas, S., & Lazcano, A. (2007). The Very Early Stages of Biological Evolution and the Nature of the Last Common Ancestor of the Three Major Cell Domains. Annual Review of Ecology, Evolution, and Systematics, 38(1), 361–379. 10.1146/annurev.ecolsys.38.091206.095825

Benler, S., Yutin, N., Antipov, D., Rayko, M., Shmakov, S., Gussow, A. B., Pevzner, P., & Koonin, E. V. (2021). Thousands of previously unknown phages discovered in whole-community human gut metagenomes. Microbiome, 9(1), 78. 10.1186/s40168-021-01017-w

Berg, M., & Roux, S. (2021). Extreme dimensions — how big (or small) can tailed phages be? Nature Reviews Microbiology 2021 19:7, 19(7), 407–407. 10.1038/s41579-021-00574-z

Brister, J. R., Ako-adjei, D., Bao, Y., & Blinkova, O. (2015). NCBI Viral Genomes Resource. Nucleic Acids Research, 43(D1), D571–D577. 10.1093/nar/gku1207

Caetano-Anollés, G. (2024). Are Viruses Taxonomic Units? A Protein Domain and Loop-Centric Phylogenomic Assessment. Viruses, 16(7), 1061. 10.3390/V16071061/S1

Caetano-Anollés, G., Wang, M., Caetano-Anollés, D., & Mittenthal, J. E. (2009). The origin, evolution and structure of the protein world. Biochemical Journal, 417(3), 621–637. 10.1042/BJ20082063

Casjens, S. R., & Gilcrease, E. B. (2009). Determining DNA Packaging Strategy by Analysis of the Termini of the Chromosomes in Tailed-Bacteriophage Virions. In Bacteriophages. Methods in Molecular Biology^TM^ (Vol. 502, pp. 91–111). 10.1007/978-1-60327-565-1_7

Caspar, D. L. D., & Klug, A. (1962). Physical Principles in the Construction of Regular Viruses. Cold Spring Harbor Symposia on Quantitative Biology, 27(0), 1–24. 10.1101/SQB.1962.027.001.005

Chen, D. H., Baker, M. L., Hryc, C. F., DiMaio, F., Jakana, J., Wu, W., Dougherty, M., Haase-Pettingell, C., Schmid, M. F., Jiang, W., Baker, D., King, J. A., & Chiu, W. (2011). Structural basis for scaffolding-mediated assembly and maturation of a dsDNA virus. Proceedings of the National Academy of Sciences of the United States of America, 108(4), 1355–1360. 10.1073/PNAS.1015739108/SUPPL_FILE/SM03.MP4

Chmelyuk, N. S., Oda, V. V., Gabashvili, A. N., & Abakumov, M. A. (2023). Encapsulins: Structure, Properties, and Biotechnological Applications. Biochemistry (Moscow) 2023 88:1, 88(1), 35–49. 10.1134/S0006297923010042

Choi, K. H., Morais, M. C., Anderson, D. L., & Rossmann, M. G. (2006). Determinants of Bacteriophage 29 Head Morphology. Structure, 14(11), 1723–1727. 10.1016/j.str.2006.09.007

Cobián Güemes, A. G., Youle, M., Cantú, V. A., Felts, B., Nulton, J., & Rohwer, F. (2016). Viruses as Winners in the Game of Life. Annual Review of Virology, 3(1), 197–214. 10.1146/annurev-virology-100114-054952

Cornell, C. E., Black, R. A., Xue, M., Litz, H. E., Ramsay, A., Gordon, M., Mileant, A., Cohen, Z. R., Williams, J. A., Lee, K. K., Drobny, G. P., & Keller, S. L. (2019). Prebiotic amino acids bind to and stabilize prebiotic fatty acid membranes. Proceedings of the National Academy of Sciences, 116(35), 17239–17244. 10.1073/pnas.1900275116

Cui, J., Schlub, T. E., & Holmes, E. C. (2014). An Allometric Relationship between the Genome Length and Virion Volume of Viruses. Journal of Virology, 88(11), 6403–6410. 10.1128/JVI.00362-14

Dai, X., & Hong Zhou, Z. (2018). Structure of the herpes simplex virus 1 capsid with associated tegument protein complexes. *Science (New York*, N.Y.), 360(6384). 10.1126/SCIENCE.AAO7298

de la Higuera, I., Kasun, G. W., Torrance, E. L., Pratt, A. A., Maluenda, A., Colombet, J., Bisseux, M., Ravet, V., Dayaram, A., Stainton, D., Kraberger, S., Zawar-Reza, P., Goldstien, S., Briskie, J. V., White, R., Taylor, H., Gomez, C., Ainley, D. G., Harding, J. S.,… Stedman, K. M. (2020). Unveiling crucivirus diversity by mining metagenomic data. MBio, 11(5), 1–17. 10.1128/MBIO.01410-20/SUPPL_FILE/MBIO.01410-20-S0001.DOCX

Dearborn, A. D., Wall, E. A., Kizziah, J. L., Klenow, L., Parker, L. K., Manning, K. A., Spilman, M. S., Spear, J. M., Christie, G. E., & Dokland, T. (2017). Competing scaffolding proteins determine capsid size during mobilization of Staphylococcus aureus pathogenicity islands. ELife, 6. 10.7554/ELIFE.30822

deCarvalho, T., Mascolo, E., Caruso, S. M., López-Pérez, J., Weston-Hafer, K., Shaffer, C., & Erill, I. (2023). Simultaneous entry as an adaptation to virulence in a novel satellite-helper system infecting Streptomyces species. The ISME Journal, 17(12), 2381–2388. 10.1038/S41396-023-01548-0

Duda, R. L., & Teschke, C. M. (2019). The amazing HK97 fold: versatile results of modest differences. Current Opinion in Virology, 36, 9–16. 10.1016/j.coviro.2019.02.001

Fillol-Salom, A., Martínez-Rubio, R., Abdulrahman, R. F., Chen, J., Davies, R., & Penadés, J. R. (2018). Phage-inducible chromosomal islands are ubiquitous within the bacterial universe. The ISME Journal, 12(9), 2114–2128. 10.1038/s41396-018-0156-3

Flamholz, Z. N., Biller, S. J., & Kelly, L. (2024). Large language models improve annotation of prokaryotic viral proteins. Nature Microbiology, 9(2), 537–549. 10.1038/s41564-023-01584-8

Fokine, A., & Rossmann, M. G. (2014). Molecular architecture of tailed double-stranded DNA phages. Bacteriophage, 4(2), e28281. 10.4161/bact.28281

Giessen, T. W. (2016). Encapsulins: microbial nanocompartments with applications in biomedicine, nanobiotechnology and materials science. Current Opinion in Chemical Biology, 34, 1–10. 10.1016/j.cbpa.2016.05.013

Giessen, T. W., Orlando, B. J., Verdegaal, A. A., Chambers, M. G., Gardener, J., Bell, D. C., Birrane, G., Liao, M., & Silver, P. A. (2019). Large protein organelles form a new iron sequestration system with high storage capacity. ELife, 8. 10.7554/eLife.46070

Giessen, T. W., & Silver, P. A. (2017). Widespread distribution of encapsulin nanocompartments reveals functional diversity. Nature Microbiology, 2(6), 17029. 10.1038/nmicrobiol.2017.29

Gilbert, W. (1986). Origin of life: The RNA world. Nature, 319(6055), 618–618. 10.1038/319618a0

Gómez-Barrera, S. N., Delgado-Tapia, W. Á., Hernández-Gutiérrez, A. E., Cayetano-Cruz, M., Méndez, C., & Bustos-Jaimes, I. (2025). Surface Engineering of the Encapsulin Nanocompartment of *Myxococcus xanthus* for Cell-Targeted Protein Delivery. ACS Omega, 10(7), 7142–7152. 10.1021/acsomega.4c10285

González, B., Monroe, L., Li, K., Yan, R., Wright, E., Walter, T., Kihara, D., Weintraub, S. T., Thomas, J. A., Serwer, P., & Jiang, W. (2020). Phage G Structure at 6.1 Å Resolution, Condensed DNA, and Host Identity Revision to a Lysinibacillus. Journal of Molecular Biology, 432(14), 4139–4153. 10.1016/j.jmb.2020.05.016

Hagberg, A., Swart, P. J., & Schult, D. A. (2008). Exploring network structure, dynamics, and function using NetworkX.

Haggård-Liungquist, E., Jacobsen, E., Rishovd, S., Six, E. W., Nilssen, Ø., Sunshine, M. G., Lindqvist, B. H., Kim, K.-J., Barreiro, V., Koonin, E. V., & Calendar, R. (1995). Bacteriophage P2: Genes Involved in Baseplate Assembly. Virology, 213(1), 109–121. 10.1006/viro.1995.1551

Hardy, J. M., Dunstan, R. A., Grinter, R., Belousoff, M. J., Wang, J., Pickard, D., Venugopal, H., Dougan, G., Lithgow, T., & Coulibaly, F. (2020). The architecture and stabilisation of flagellotropic tailed bacteriophages. Nature Communications, 11(1), 3748. 10.1038/s41467-020-17505-w

Hawkins, D. E. D. P., Bayfield, O. W., Fung, H. K. H., Grba, D. N., Huet, A., Conway, J. F., & Antson, A. A. (2023). Insights into a viral motor: the structure of the HK97 packaging termination assembly. Nucleic Acids Research, 51(13), 7025. 10.1093/NAR/GKAD480

Helgstrand, C., Wikoff, W. R., Duda, R. L., Hendrix, R. W., Johnson, J. E., & Liljas, L. (2003). The Refined Structure of a Protein Catenane: The HK97 Bacteriophage Capsid at 3.44 Å Resolution. Journal of Molecular Biology, 334(5), 885–899. 10.1016/J.JMB.2003.09.035

Hendrix, R. W. (2005). Bacteriophage HK97: Assembly of the Capsid and Evolutionary Connections. In Advances in Virus Research (Vol. 64, pp. 1–14). Academic Press. 10.1016/S0065-3527(05)64001-8

Heymann, J. B., Cheng, N., Newcomb, W. W., Trus, B. L., Brown, J. C., & Steven, A. C. (2003). Dynamics of herpes simplex virus capsid maturation visualized by time-lapse cryo-electron microscopy. Nature Structural & Molecular Biology, 10(5), 334–341. 10.1038/nsb922

Howard-Varona, C., Hargreaves, K. R., Abedon, S. T., & Sullivan, M. B. (2017). Lysogeny in nature: mechanisms, impact and ecology of temperate phages. The ISME Journal, 1–10. 10.1038/ismej.2017.16

Hryc, C. F., Chen, D.-H., Afonine, P. V., Jakana, J., Wang, Z., Haase-Pettingell, C., Jiang, W., Adams, P. D., King, J. A., Schmid, M. F., & Chiu, W. (2017). Accurate model annotation of a near-atomic resolution cryo-EM map. Proceedings of the National Academy of Sciences, 114(12), 3103–3108. 10.1073/pnas.1621152114

Hua, J., Huet, A., Lopez, C. A., Toropova, K., Pope, W. H., Duda, R. L., Hendrix, R. W., & Conway, J. F. (2017). Capsids and Genomes of Jumbo-Sized Bacteriophages Reveal the Evolutionary Reach of the HK97 Fold. MBio, 8(5). 10.1128/mBio.01579-17

Huet, A., Oh, B., Maurer, J., Duda, R. L., & Conway, J. F. (2023). A symmetry mismatch unraveled: How phage HK97 scaffold flexibly accommodates a 12-fold pore at a 5-fold viral capsid vertex. Science Advances, 9(24). 10.1126/sciadv.adg8868

Hyatt, D., Chen, G.-L., LoCascio, P. F., Land, M. L., Larimer, F. W., & Hauser, L. J. (2010). Prodigal: prokaryotic gene recognition and translation initiation site identification. BMC Bioinformatics, 11(1), 119. 10.1186/1471-2105-11-119

Ibarra, B., Castón, J. R., Llorca, O., Valle, M., Valpuesta, J. M., & Carrascosa, J. L. (2000). Topology of the components of the DNA packaging machinery in the phage φ29 prohead. Journal of Molecular Biology, 298(5), 807–815. 10.1006/jmbi.2000.3712

Ignatiou, A., Brasilès, S., El Sadek Fadel, M., Bürger, J., Mielke, T., Topf, M., Tavares, P., & Orlova, E. V. (2019). Structural transitions during the scaffolding-driven assembly of a viral capsid. Nature Communications 2019 10:1, 10(1), 1–11. 10.1038/s41467-019-12790-6

Ionel, A., Velázquez-Muriel, J. A., Luque, D., Cuervo, A., Castón, J. R., Valpuesta, J. M., Martín-Benito, J., & Carrascosa, J. L. (2011). Molecular Rearrangements Involved in the Capsid Shell Maturation of Bacteriophage T7. Journal of Biological Chemistry, 286(1), 234–242. 10.1074/jbc.M110.187211

Ivanova, N., Tringe, S. G., Liolios, K., Liu, W., Morrison, N., Hugenholtz, P., & Kyrpides, N. C. (2010). A call for standardized classification of metagenome projects. Environmental Microbiology, 12(7), 1803–1805. 10.1111/j.1462-2920.2010.02270.x

Jones, J. A., Andreas, M. P., & Giessen, T. W. (2024). Structural basis for peroxidase encapsulation inside the encapsulin from the Gram-negative pathogen Klebsiella pneumoniae. Nature Communications, 15(1), 2558. 10.1038/s41467-024-46880-x

Jumper, J., Evans, R., Pritzel, A., Green, T., Figurnov, M., Ronneberger, O., Tunyasuvunakool, K., Bates, R., Žídek, A., Potapenko, A., Bridgland, A., Meyer, C., Kohl, S. A. A., Ballard, A. J., Cowie, A., Romera-Paredes, B., Nikolov, S., Jain, R., Adler, J.,… Hassabis, D. (2021). Highly accurate protein structure prediction with AlphaFold. Nature, 596(7873), 583–589. 10.1038/s41586-021-03819-2

Kashif-Khan, N., Savva, R., & Frank, S. (2024). Mining metagenomics data for novel bacterial nanocompartments. NAR Genomics and Bioinformatics, 6(1). 10.1093/nargab/lqae025

Katoh, K. (2002). MAFFT: a novel method for rapid multiple sequence alignment based on fast Fourier transform. Nucleic Acids Research, 30(14), 3059–3066. 10.1093/nar/gkf436

Katoh, K., & Toh, H. (2008). Recent developments in the MAFFT multiple sequence alignment program. Briefings in Bioinformatics, 9(4), 286–298. 10.1093/bib/bbn013

Kauffman, S. A. (1971). Cellular Homeostasis, Epigenesis and Replication in Randomly Aggregated Macromolecular Systems. Journal of Cybernetics, 1(1), 71–96. 10.1080/01969727108545830

Kazlauskas, D., Varsani, A., Koonin, E. V., & Krupovic, M. (2019). Multiple origins of prokaryotic and eukaryotic single-stranded DNA viruses from bacterial and archaeal plasmids. Nature Communications, 10(1), 3425. 10.1038/s41467-019-11433-0

Kejzar, N., Laanto, E., Rissanen, I., Abrishami, V., Selvaraj, M., Moineau, S., Ravantti, J., Sundberg, L. R., & Huiskonen, J. T. (2022). Cryo-EM structure of ssDNA bacteriophage ΦCjT23 provides insight into early virus evolution. Nature Communications 2022 13:1, 13(1), 1–13. 10.1038/s41467-022-35123-6

Kieft, K., Zhou, Z., & Anantharaman, K. (2020). VIBRANT: automated recovery, annotation and curation of microbial viruses, and evaluation of viral community function from genomic sequences. Microbiome, 8(1), 90. 10.1186/s40168-020-00867-0

Kirchberger, P. C., & Ochman, H. (2023). Microviruses: A World Beyond φX174. Annual Review of Virology, 10(1), 99–118. 10.1146/ANNUREV-VIROLOGY-100120-011239/CITE/REFWORKS

Kizziah, J. L., Rodenburg, C. M., & Dokland, T. (2020). Structure of the Capsid Size-Determining Scaffold of “Satellite” Bacteriophage P4. Viruses, 12(9), 953. 10.3390/v12090953

Koonin, E. V., Dolja, V. V., Krupovic, M., Varsani, A., Wolf, Y. I., Yutin, N., Zerbini, F. M., & Kuhn, J. H. (2020). Global Organization and Proposed Megataxonomy of the Virus World. Microbiology and Molecular Biology Reviews, 84(2). 10.1128/MMBR.00061-19

Koonin, E. V, Kuhn, J. H., Dolja, V. V, & Krupovic, M. (2024). Megataxonomy and global ecology of the virosphere. The ISME Journal, 18(1). 10.1093/ismejo/wrad042

Krupovic, M., Dolja, V. V., & Koonin, E. V. (2019). Origin of viruses: primordial replicators recruiting capsids from hosts. Nature Reviews Microbiology, 17(7), 449–458. 10.1038/s41579-019-0205-6

Krupovic, M., & Koonin, E. V. (2017). Multiple origins of viral capsid proteins from cellular ancestors. Proceedings of the National Academy of Sciences, 114(12), E2401–E2410. 10.1073/pnas.1621061114

Krupovic, M., Kuhn, J. H., Fischer, M. G., & Koonin, E. V. (2024). Natural history of eukaryotic DNA viruses with double jelly-roll major capsid proteins. Proceedings of the National Academy of Sciences of the United States of America, 121(23), e2405771121. 10.1073/PNAS.2405771121/SUPPL_FILE/PNAS.2405771121.SAPP.PDF

Laanto, E., Mäntynen, S., De Colibus, L., Marjakangas, J., Gillum, A., Stuart, D. I., Ravantti, J. J., Huiskonen, J. T., & Sundberg, L. R. (2017). Virus found in a boreal lake links ssDNA and dsDNA viruses. Proceedings of the National Academy of Sciences of the United States of America, 114(31), 8378–8383. 10.1073/PNAS.1703834114/SUPPL_FILE/PNAS.201703834SI.PDF

Lander, G. C., Johnson, J. E., Rau, D. C., Potter, C. S., Carragher, B., & Evilevitch, A. (2013). DNA bending-induced phase transition of encapsidated genome in phage λ. Nucleic Acids Research, 41(8), 4518–4524. 10.1093/NAR/GKT137

Laurenceau, R., Raho, N., Forget, M., Arellano, A. A., & Chisholm, S. W. (2020). Frequency of mispackaging of Prochlorococcus DNA by cyanophage. The ISME Journal 2020 15:1, 15(1), 129–140. 10.1038/s41396-020-00766-0

Lee, D. Y., Bartels, C., McNair, K., Edwards, R. A., Swairjo, M. A., & Luque, A. (2022). Predicting the capsid architecture of phages from metagenomic data. Computational and Structural Biotechnology Journal, 20, 721–732. 10.1016/j.csbj.2021.12.032

Letunic, I., & Bork, P. (2024). Interactive Tree of Life (iTOL) v6: recent updates to the phylogenetic tree display and annotation tool. Nucleic Acids Research, 52(W1), W78–W82. 10.1093/nar/gkae268

Luque, A., Benler, S., Lee, D. Y., Brown, C., & White, S. (2020). The Missing Tailed Phages: Prediction of Small Capsid Candidates. Microorganisms, 8(12), 1944. 10.3390/microorganisms8121944

Luque, A., & Reguera, D. (2010). The Structure of Elongated Viral Capsids. Biophysical Journal, 98(12), 2993–3003. 10.1016/j.bpj.2010.02.051

Luque, A., & Silveira, C. B. (2020). Quantification of lysogeny caused by phage coinfections in microbial communities from biophysical principles. MSystems, 5, e00353–20. 10.1128/mSystems.00353-20

Luque, A., Zandi, R., & Reguera, D. (2010). Optimal architectures of elongated viruses. PNAS, 107(12), 1–6. 10.1073/pnas.0915122107

Mankertz, A., & Hillenbrand, B. (2001). Replication of porcine circovirus type 1 requires two proteins encoded by the viral rep gene. Virology, 279(2), 429–438. 10.1006/VIRO.2000.0730

Mansy, S. S., & Szostak, J. W. (2009). Reconstructing the Emergence of Cellular Life through the Synthesis of Model Protocells. Cold Spring Harbor Symposia on Quantitative Biology, 74(0), 47–54. 10.1101/sqb.2009.74.014

Medvedeva, S., Sun, J., Yutin, N., Koonin, E. V., Nunoura, T., Rinke, C., & Krupovic, M. (2022). Three families of Asgard archaeal viruses identified in metagenome-assembled genomes. Nature Microbiology 2022 7:7, 7(7), 962–973. 10.1038/s41564-022-01144-6

Meng, E. C., Goddard, T. D., Pettersen, E. F., Couch, G. S., Pearson, Z. J., Morris, J. H., & Ferrin, T. E. (2023). <scp>UCSF ChimeraX</scp>: Tools for structure building and analysis. Protein Science, 32(11), e4792. 10.1002/pro.4792

Minh, B. Q., Schmidt, H. A., Chernomor, O., Schrempf, D., Woodhams, M. D., von Haeseler, A., & Lanfear, R. (2020). IQ-TREE 2: New Models and Efficient Methods for Phylogenetic Inference in the Genomic Era. Molecular Biology and Evolution, 37(5), 1530–1534. 10.1093/molbev/msaa015

Mistry, J., Chuguransky, S., Williams, L., Qureshi, M., Salazar, G. A., Sonnhammer, E. L. L., Tosatto, S. C. E., Paladin, L., Raj, S., Richardson, L. J., Finn, R. D., & Bateman, A. (2021). Pfam: The protein families database in 2021. Nucleic Acids Research, 49(D1), D412–D419. 10.1093/nar/gkaa913

Moi, D., Bernard, C., Steinegger, M., Nevers, Y., Langleib, M., & Dessimoz, C. (2023). Structural phylogenetics unravels the evolutionary diversification of communication systems in gram-positive bacteria and their viruses. BioRxiv, 2023.09.19.558401. 10.1101/2023.09.19.558401

Morais, M. C., Kanamaru, S., Badasso, M. O., Koti, J. S., Owen, B. A. L., McMurray, C. T., Anderson, D. L., & Rossmann, M. G. (2003). Bacteriophage φ29 scaffolding protein gp7 before and after prohead assembly. Nature Structural & Molecular Biology, 10(7), 572–576. 10.1038/nsb939

Moreira, D., & López-García, P. (2009). Ten reasons to exclude viruses from the tree of life. Nature Reviews Microbiology 2009 7:4, 7(4), 306–311. 10.1038/nrmicro2108

Mutti, G., Ocaña-Pallarès, E., & Gabaldón, T. (2025). Newly Developed Structure-Based Methods Do Not Outperform Standard Sequence-Based Methods for Large-Scale Phylogenomics. Molecular Biology and Evolution, 42(7). 10.1093/molbev/msaf149

Nasir, A., & Caetano-Anollés, G. (2015). A phylogenomic data-driven exploration of viral origins and evolution. Science Advances, 1(8). 10.1126/SCIADV.1500527/SUPPL_FILE/NASIRTABLES7.XLSX

Nayfach, S., Camargo, A. P., Schulz, F., Eloe-Fadrosh, E., Roux, S., & Kyrpides, N. C. (2021). CheckV assesses the quality and completeness of metagenome-assembled viral genomes. Nature Biotechnology, 39(5), 578–585. 10.1038/s41587-020-00774-7

Nayfach, S., Shi, Z. J., Seshadri, R., Pollard, K. S., & Kyrpides, N. C. (2019). New insights from uncultivated genomes of the global human gut microbiome. Nature, 568(7753), 505–510. 10.1038/s41586-019-1058-x

Nichols, R. J., LaFrance, B., Phillips, N. R., Radford, D. R., Oltrogge, L. M., Valentin-Alvarado, L. E., Bischoff, A. J., Nogales, E., & Savage, D. F. (2021). Discovery and characterization of a novel family of prokaryotic nanocompartments involved in sulfur metabolism. ELife, 10. 10.7554/eLife.59288

Oh, B., Moyer, C. L., Hendrix, R. W., & Duda, R. L. (2014). The delta domain of the HK97 major capsid protein is essential for assembly. Virology, 456–457(1), 171–178. 10.1016/J.VIROL.2014.03.022

Olsen, N. S., Hendriksen, N. B., Hansen, L. H., & Kot, W. (2020). A New High-Throughput Screening Method for Phages: Enabling Crude Isolation and Fast Identification of Diverse Phages with Therapeutic Potential. PHAGE, 1(3), 137–148. 10.1089/phage.2020.0016

Paez-Espino, D., Eloe-Fadrosh, E. A., Pavlopoulos, G. A., Thomas, A. D., Huntemann, M., Mikhailova, N., Rubin, E., Ivanova, N. N., & Kyrpides, N. C. (2016). Uncovering Earth’s virome. Nature, 536(7617), 425–430. 10.1038/nature19094

Paez-Espino, D., Roux, S., Chen, I.-M. A., Palaniappan, K., Ratner, A., Chu, K., Huntemann, M., Reddy, T. B. K., Pons, J. C., Llabrés, M., Eloe-Fadrosh, E. A., Ivanova, N. N., & Kyrpides, N. C. (2019). IMG/VR v.2.0: an integrated data management and analysis system for cultivated and environmental viral genomes. Nucleic Acids Research, 47(D1), D678–D686. 10.1093/nar/gky1127

Pasolli, E., Asnicar, F., Manara, S., Zolfo, M., Karcher, N., Armanini, F., Beghini, F., Manghi, P., Tett, A., Ghensi, P., Collado, M. C., Rice, B. L., DuLong, C., Morgan, X. C., Golden, C. D., Quince, C., Huttenhower, C., & Segata, N. (2019). Extensive Unexplored Human Microbiome Diversity Revealed by Over 150,000 Genomes from Metagenomes Spanning Age, Geography, and Lifestyle. Cell, 176(3), 649–662.e20. 10.1016/j.cell.2019.01.001

Penadés, J. R., & Christie, G. E. (2015). The Phage-Inducible Chromosomal Islands: A Family of Highly Evolved Molecular Parasites. Annual Review of Virology, 2(1), 181–201. 10.1146/annurev-virology-031413-085446

Penadés, J. R., Seed, K. D., Chen, J., Bikard, D., & Rocha, E. P. C. (2025). Genetics, ecology and evolution of phage satellites. Nature Reviews Microbiology 2025, 1–13. 10.1038/s41579-025-01156-z

Pietilä, M. K., Laurinmäki, P., Russell, D. A., Ko, C.-C., Jacobs-Sera, D., Hendrix, R. W., Bamford, D. H., & Butcher, S. J. (2013). Structure of the archaeal head-tailed virus HSTV-1 completes the HK97 fold story. Proceedings of the National Academy of Sciences of the United States of America, 110(26), 10604–10609. 10.1073/pnas.1303047110

Podgorski, J. M., Podgorski, J., Abad, L., Jacobs-Sera, D., Freeman, K. G., Brown, C., Hatfull, G., Luque, A., & White, S. J. (2023). A novel stabilization mechanism accommodating genome length variation in evolutionarily related viral capsids. In bioRxiv (p. 2023.11.03.565530). Cold Spring Harbor Laboratory. 10.1101/2023.11.03.565530

Pourcel, C., Essoh, C., Ouldali, M., & Tavares, P. (2024). Acinetobacter baumannii satellite phage Aci01-2-Phanie depends on a helper myophage for its multiplication. Journal of Virology, 98(7). 10.1128/JVI.00667-24/SUPPL_FILE/JVI.00667-24-S0002.XLSX

Preiner, M., Asche, S., Becker, S., Betts, H. C., Boniface, A., Camprubi, E., Chandru, K., Erastova, V., Garg, S. G., Khawaja, N., Kostyrka, G., Machné, R., Moggioli, G., Muchowska, K. B., Neukirchen, S., Peter, B., Pichlhöfer, E., Radványi, Á., Rossetto, D.,… Xavier, J. C. (2020). The Future of Origin of Life Research: Bridging Decades-Old Divisions. Life, 10(3), 20. 10.3390/life10030020

Pressman, A., Blanco, C., & Chen, I. A. (2015). The RNA World as a Model System to Study the Origin of Life. Current Biology, 25(19), R953–R963. 10.1016/j.cub.2015.06.016

Rao, V. B., Fokine, A., & Fang, Q. (2021). The Remarkable Vial Portal Vertex: structure and a plausible model for mechanism. Current Opinion in Virology, 51, 65. 10.1016/J.COVIRO.2021.09.004

Ravin, N. V. (2015). Replication and Maintenance of Linear Phage-Plasmid N15. Microbiology Spectrum, 3(1). 10.1128/MICROBIOLSPEC.PLAS-0032-2014/ASSET/DBF1500D-63E0-4CD3-B333-09FE3E88661E/ASSETS/GRAPHIC/PLAS-0032-2014-FIG5.GIF

Roos, W. H., Gertsman, I., May, E. R., Brooks, C. L., Johnson, J. E., & Wuite, G. J. L. (2012). Mechanics of bacteriophage maturation. Proceedings of the National Academy of Sciences of the United States of America, 109(7), 2342–2347. 10.1073/PNAS.1109590109/SUPPL_FILE/PNAS.1109590109_SI.PDF

Ross, J., McIver, Z., Lambert, T., Piergentili, C., Bird, J. E., Gallagher, K. J., Cruickshank, F. L., James, P., Zarazúa-Arvizu, E., Horsfall, L. E., Waldron, K. J., Wilson, M. D., Logan Mackay, C., Baslé, A., Clarke, D. J., & Marles-Wright, J. (2022). Pore dynamics and asymmetric cargo loading in an encapsulin nanocompartment. Science Advances, 8(4), 4461. 10.1126/SCIADV.ABJ4461/SUPPL_FILE/SCIADV.ABJ4461_MOVIES_S1_TO_S3.ZIP

San Martín, C., & Van Raaij, M. J. (2018). The so far farthest reaches of the double jelly roll capsid protein fold 06 Biological Sciences 0601 Biochemistry and Cell Biology. Virology Journal, 15(1), 1–6. 10.1186/S12985-018-1097-1/TABLES/1

Segré, D., Ben-Eli, D., Deamer, D. W., & Lancet, D. (2001). The Lipid World. Origins of Life and Evolution of the Biosphere, 31(1–2), 119–145. 10.1023/A:1006746807104

Selivanovitch, E., LaFrance, B., & Douglas, T. (2021). Molecular exclusion limits for diffusion across a porous capsid. Nature Communications, 12(1), 2903. 10.1038/s41467-021-23200-1

Smith, E., & Morowitz, H. J. (2004). Universality in intermediary metabolism. Proceedings of the National Academy of Sciences, 101(36), 13168–13173. 10.1073/pnas.0404922101

Söding, J. (2005). Protein homology detection by HMM–HMM comparison. Bioinformatics, 21(7), 951–960. 10.1093/bioinformatics/bti125

Sonntag, F., Köther, K., Schmidt, K., Weghofer, M., Raupp, C., Nieto, K., Kuck, A., Gerlach, B., Böttcher, B., Müller, O. J., Lux, K., Hörer, M., & Kleinschmidt, J. A. (2011). The Assembly-Activating Protein Promotes Capsid Assembly of Different Adeno-Associated Virus Serotypes. Journal of Virology, 85(23), 12686–12697. 10.1128/JVI.05359-11/ASSET/9822EA7F-6689-4755-9978-884B6872AD15/ASSETS/GRAPHIC/ZJV9990952780010.JPEG

Soto-Perez, P., Bisanz, J. E., Berry, J. D., Lam, K. N., Bondy-Denomy, J., & Turnbaugh, P. J. (2019). CRISPR-Cas System of a Prevalent Human Gut Bacterium Reveals Hyper-targeting against Phages in a Human Virome Catalog. Cell Host & Microbe, 26(3), 325–335.e5. 10.1016/j.chom.2019.08.008

Stone, N. P., Demo, G., Agnello, E., & Kelch, B. A. (2019). Principles for enhancing virus capsid capacity and stability from a thermophilic virus capsid structure. Nature Communications, 10(1), 4471. 10.1038/s41467-019-12341-z

Subramanian, S., Kerns, H. R., Braverman, S. G., & Doore, S. M. (2025). The structure of Shigella virus Sf14 reveals the presence of two decoration proteins and two long tail fibers. Communications Biology 2025 8:1, 8(1), 1–11. 10.1038/s42003-025-07668-x

Suhanovsky, M. M., & Teschke, C. M. (2015). Nature’s favorite building block: Deciphering folding and capsid assembly of proteins with the HK97-fold. Virology, 479–480, 487–497. 10.1016/j.virol.2015.02.055

Sun, L., Rossmann, M. G., & Fane, B. A. (2014). High-Resolution Structure of a Virally Encoded DNA-Translocating Conduit and the Mechanism of DNA Penetration. Journal of Virology, 88(18), 10276–10279. 10.1128/JVI.00291-14/ASSET/3DA09E36-6386-4204-B3B4-0165A20E9AD5/ASSETS/GRAPHIC/ZJV9990994710001.JPEG

Sun, L., Young, L. N., Zhang, X., Boudko, S. P., Fokine, A., Zbornik, E., Roznowski, A. P., Molineux, I. J., Rossmann, M. G., & Fane, B. A. (2013). Icosahedral bacteriophage ΦX174 forms a tail for DNA transport during infection. Nature 2013 505:7483, 505(7483), 432–435. 10.1038/nature12816

Sutter, M., Boehringer, D., Gutmann, S., Günther, S., Prangishvili, D., Loessner, M. J., Stetter, K. O., Weber-Ban, E., & Ban, N. (2008). Structural basis of enzyme encapsulation into a bacterial nanocompartment. Nature Structural & Molecular Biology, 15(9), 939–947. 10.1038/nsmb.1473

Takagi, Y. A., Nguyen, D. H., Wexler, T. B., & Goldman, A. D. (2020). The Coevolution of Cellularity and Metabolism Following the Origin of Life. Journal of Molecular Evolution, 88(7), 598–617. 10.1007/S00239-020-09961-1/FIGURES/12

Terzian, P., Olo Ndela, E., Galiez, C., Lossouarn, J., Pérez Bucio, R. E., Mom, R., Toussaint, A., Petit, M.-A., & Enault, F. (2021). PHROG: families of prokaryotic virus proteins clustered using remote homology. NAR Genomics and Bioinformatics, 3(3). 10.1093/nargab/lqab067

Twarock, R., & Luque, A. (2019). Structural puzzles in virology solved with an overarching icosahedral design principle. Nature Communications, 10(1), 4414. 10.1038/s41467-019-12367-3

van Kempen, M., Kim, S. S., Tumescheit, C., Mirdita, M., Lee, J., Gilchrist, C. L. M., Söding, J., & Steinegger, M. (2024). Fast and accurate protein structure search with Foldseek. Nature Biotechnology, 42(2), 243–246. 10.1038/s41587-023-01773-0

Veesler, D., & Johnson, J. E. (2012). Virus maturation. Annual Review of Biophysics, 41(1), 473–496. 10.1146/ANNUREV-BIOPHYS-042910-155407/CITE/REFWORKS

Vincent, L., Berg, M., Krismer, M., Saghafi, S. T., Cosby, J., Sankari, T., Vetsigian, K., Cleaves, H. J., & Baum, D. A. (2019). Chemical Ecosystem Selection on Mineral Surfaces Reveals Long-Term Dynamics Consistent with the Spontaneous Emergence of Mutual Catalysis. Life, 9(4), 80. 10.3390/life9040080

Wächtershäuser, G. (1997). The Origin of Life and its Methodological Challenge. Journal of Theoretical Biology, 187(4), 483–494. 10.1006/jtbi.1996.0383

Wang, Y., & Douglas, T. (2022). Bioinspired Approaches to Self-Assembly of Virus-like Particles: From Molecules to Materials. Accounts of Chemical Research, 55(10), 1349– 1359. 10.1021/acs.accounts.2c00056

Wigington, C. H., Sonderegger, D., Brussaard, C. P. D., Buchan, A., Finke, J. F., Fuhrman, J. A., Lennon, J. T., Middelboe, M., Suttle, C. A., Stock, C., Wilson, W. H., Wommack, K. E., Wilhelm, S. W., & Weitz, J. S. (2016). Re-examination of the relationship between marine virus and microbial cell abundances. Nature Microbiology, 1(3), 15024. 10.1038/nmicrobiol.2015.24

Wikoff, W. R., Conway, J. F., Tang, J., Lee, K. K., Gan, L., Cheng, N., Duda, R. L., Hendrix, R. W., Steven, A. C., & Johnson, J. E. (2006). Time-resolved molecular dynamics of bacteriophage HK97 capsid maturation interpreted by electron cryo-microscopy and X-ray crystallography. Journal of Structural Biology, 153(3), 300–306. 10.1016/J.JSB.2005.11.009

Wikoff, W. R., Liljas, L., Duda, R. L., Tsuruta, H., Hendrix, R. W., & Johnson, J. E. (2000). Topologically Linked Protein Rings in the Bacteriophage HK97 Capsid. Science, 289(5487), 2129–2133. 10.1126/SCIENCE.289.5487.2129

Williams, E. M., Jung, S. M., Coffman, J. L., & Lutz, S. (2018). Pore Engineering for Enhanced Mass Transport in Encapsulin Nanocompartments. ACS Synthetic Biology, 7(11), 2514– 2517. 10.1021/ACSSYNBIO.8B00295

Wommack, K. E., & Colwell, R. R. (2000). Virioplankton: Viruses in Aquatic Ecosystems. Microbiology and Molecular Biology Reviews, 64(1), 69–114. 10.1128/MMBR.64.1.69-114.2000

Woodson, M., Pajak, J., Mahler, B. P., Zhao, W., Zhang, W., Arya, G., White, M. A., Jardine, P. J., & Morais, M. C. (2021). A viral genome packaging motor transitions between cyclic and helical symmetry to translocate dsDNA. Science Advances, 7(19). 10.1126/sciadv.abc1955

Wu, W., Newcomb, W. W., Cheng, N., Aksyuk, A., Winkler, D. C., & Steven, A. C. (2016). Internal Proteins of the Procapsid and Mature Capsids of Herpes Simplex Virus 1 Mapped by Bubblegram Imaging. Journal of Virology, 90(10), 5176–5186. 10.1128/JVI.03224-15

Xu, J., Wang, D., Gui, M., & Xiang, Y. (2019). Structural assembly of the tailed bacteriophage 29. Nature Communications 2019 10:1, 10(1), 1–16. 10.1038/s41467-019-10272-3

Yap, M. L., & Rossmann, M. G. (2014). Structure and function of bacteriophage T4. Future Microbiology, 9(12), 1319–1337. 10.2217/FMB.14.91

