## Supplementary Materials for "Small viruses reveal bidirectional evolution between HK97-fold viruses and encapsulins via procapsids" (from another version of this preprint: version 2, 2025-08-28)

This file contains all the supplementary materials for the manuscript titled "Small uncultured viral-like entities redraw the origin of viruses and cellular compartments" by Aguilar *et al*. Supplementary materials are divided into two sections: 1) Supplementary Figures, and 2) Links to data files.

### Section 1. Supplementary figures

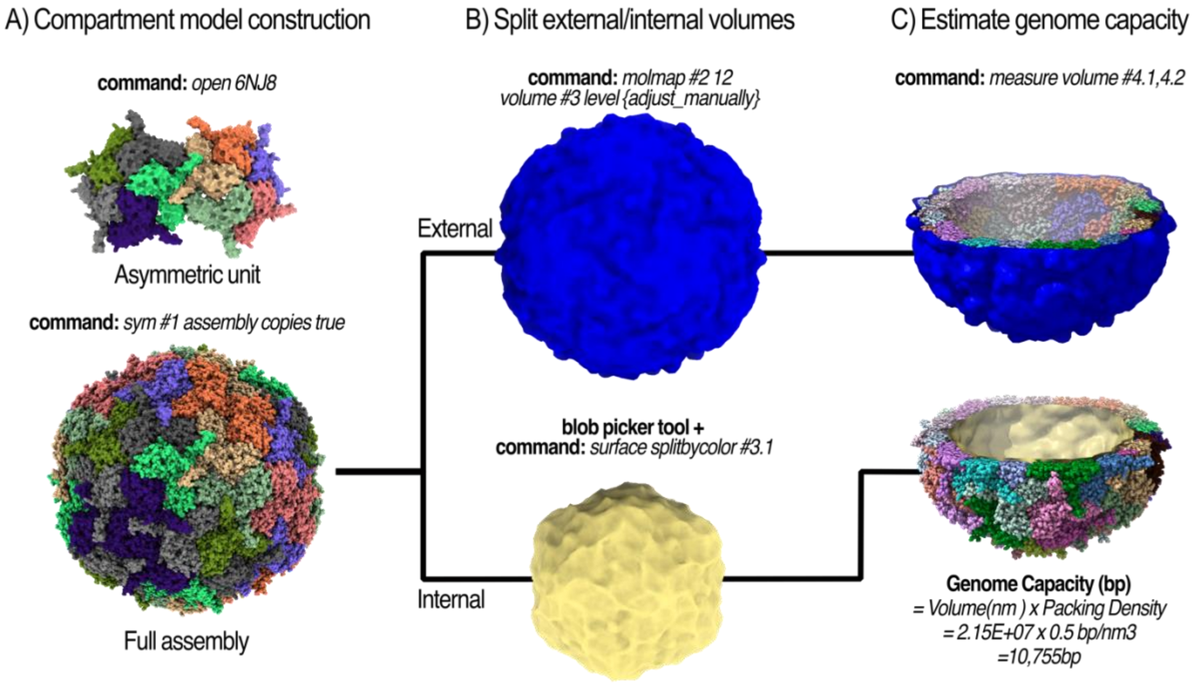


**Figure S1.** Workflow for measuring the enclosed volume of HK97-fold compartments using ChimeraX. A) Commands used to reconstruct the full capsid assembly from asymmetric unit coordinates, generating the complete surface model. B) Commands to create a molecular density map from the surface model, split the external and internal surfaces, and isolate the internal cavity for volume analysis. C) Measurement of the enclosed internal volume using ChimeraX's measure volume command, and projection of genome capacity based on the estimated internal volume.

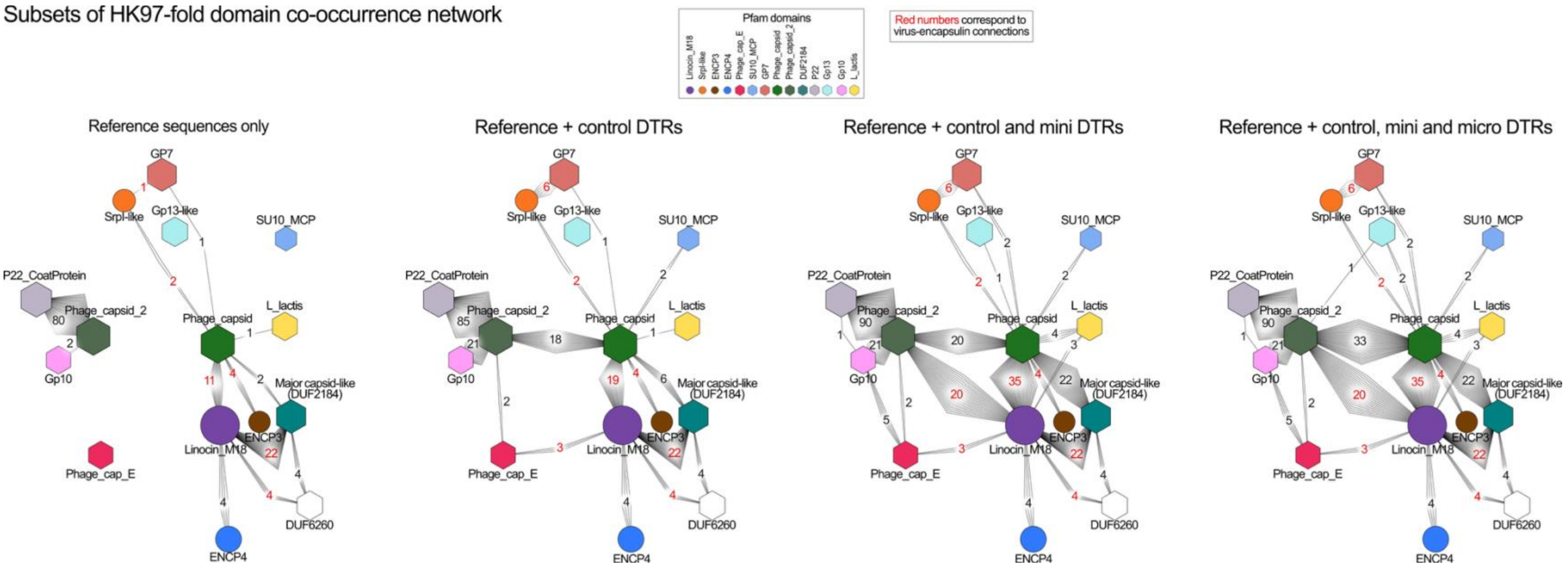


**Figure S2.** Co-occurrence network of domains with different subsets. Nodes represent individual Pfam profiles, node size scales with total degree. Edges link profiles found in the same protein, with edge labels showing co-occurrence counts (red numbers denote counts of non-canonical virus–encapsulin links).

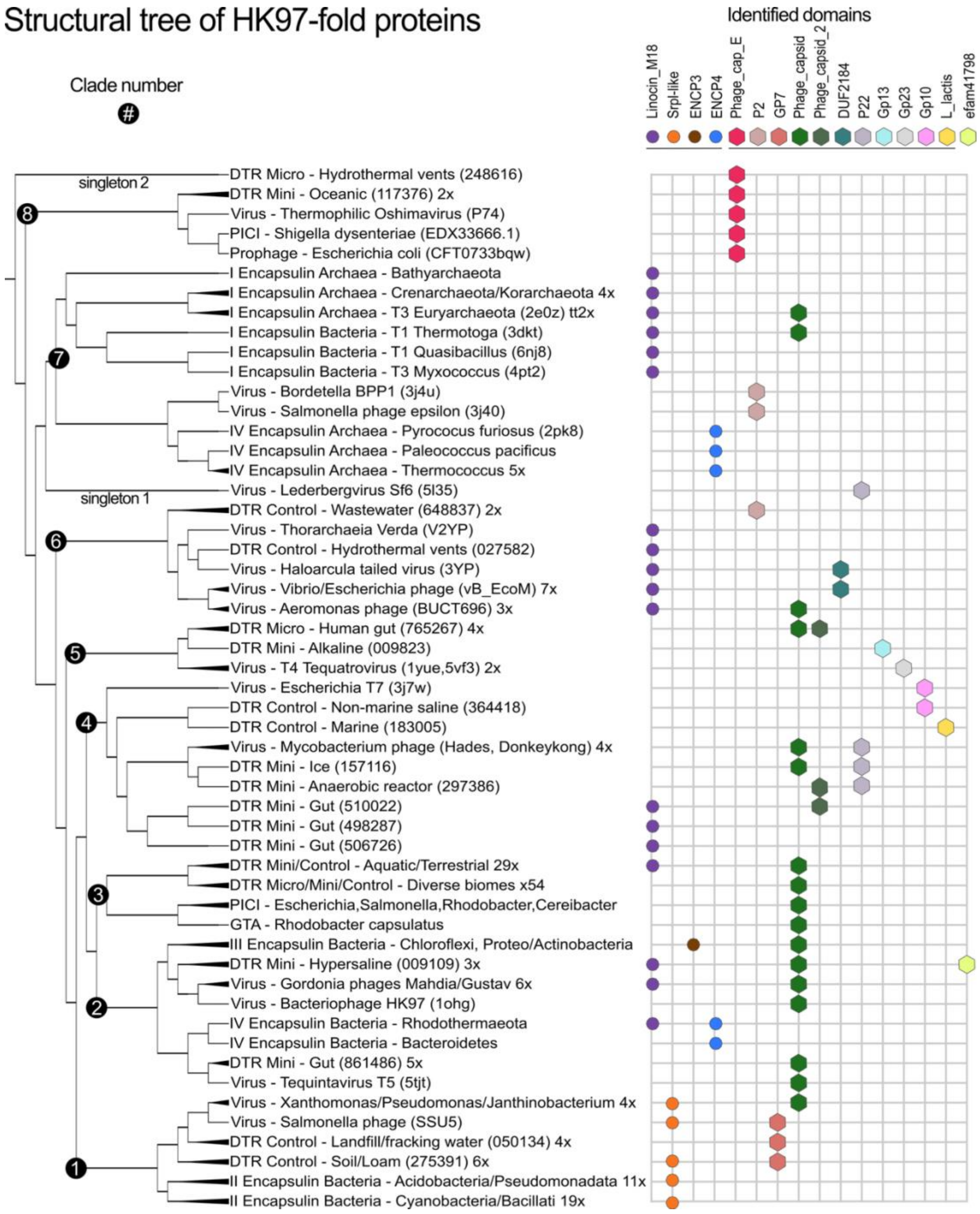


**Figure S3.** Complete rooted structural tree. The tree compares HK97-fold structures, including encapsulins, twilight-zone viral MCPs, and RefSeq representatives. The number of each clade and singleton is indicated. Colapsed Each leaf includes the associated domains from the *phage_coat* clan to the right, based on the color coding displayed in the legend on top.

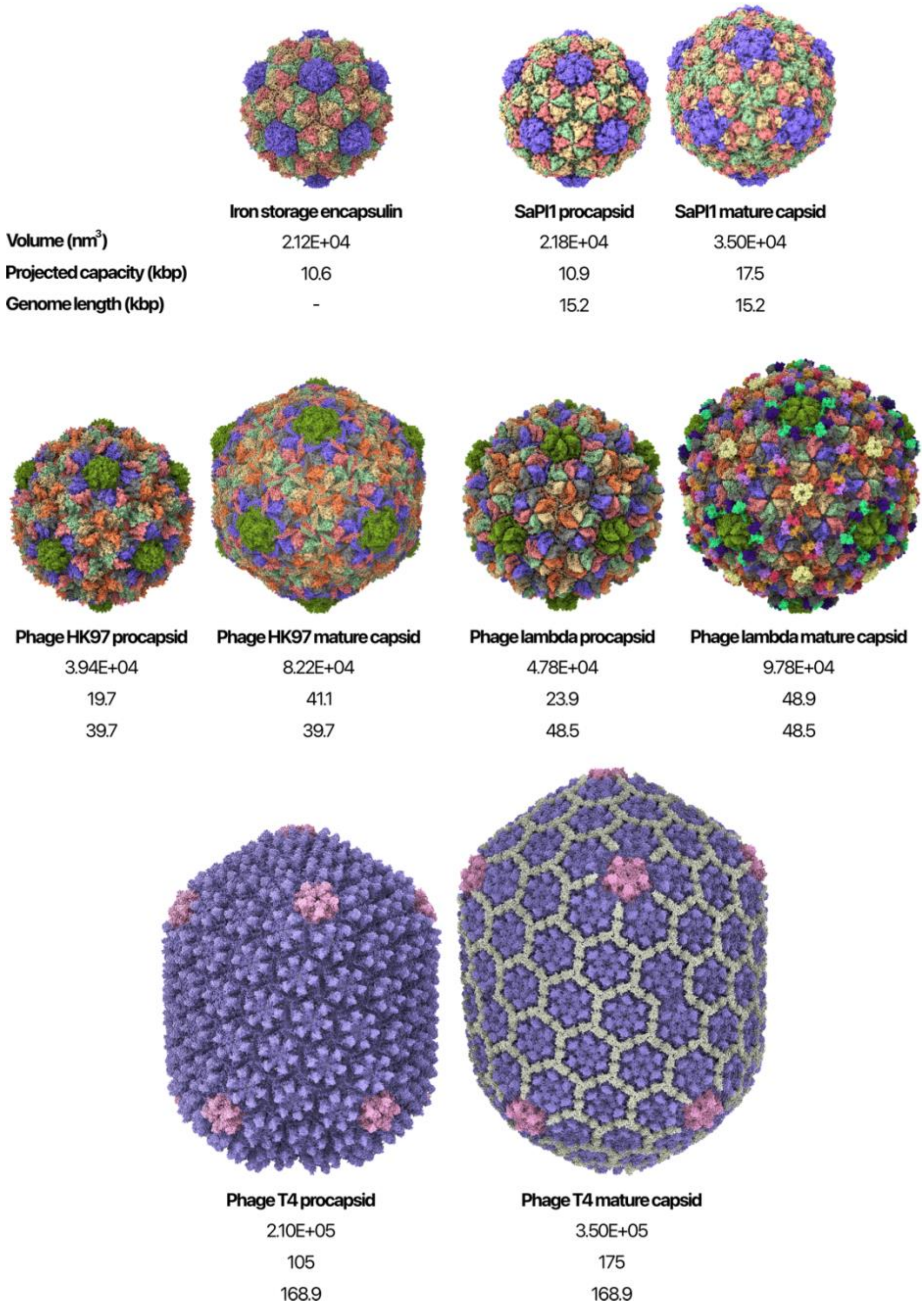


**Figure S4**. Structural comparison of encapsulin and viral capsid architectures across different triangulation numbers (T). The figure shows atomic reconstructions of an encapsulin (T=4; iron storage encapsulin PDB: 6NJ8), procapsid and mature capsid of Staphylococcus aureus Pathogenicity Island 1 (SaPI1, T=4; procapsid: PDB 7RWZ, mature capsid: PDB 8VD4), Enterobacteria phage HK97 (T=7; procapsid: PDB 3E8K, mature capsid: PDB 1OHG), Enterobacteria phage λ (T=7; procapsid: PDB 7VI9, mature capsid: PDB 7VII), and Enterobacteria phage T4 (T=13 prolate; procapsid: PDB 7VRT, mature capsid: PDB 7VS5).

### Section 2. Links to Data Files

All files are available for download at:
https://drive.google.com/drive/folders/1s4JYU9VXwNfHClpkb8KhwEveDSAKeLWu?usp=sharing

##### *Environmental viral genomes*

`data_file_S1_IMG_datasets.xlsx`
Complete list of Joint Genome Institute, Integrated Microbial Genomes & Microbiomes associated datasets. This file contains the identifiers of each study, as well as its associated DOIs.

`data_file_S2_biomes.csv`
Hierarchical biome classification schema used to map metagenomic contigs to ecological categories, following Ivanova et al. (2010). Includes all possible biome categories at multiple levels.

`data_file_S3_associated_biomes.csv`
Metadata table links each environmental viral contig to its assigned biome(s), along with relevant sample-level information.

`data_file_S4_genomes.fna`
FASTA file containing all environmental viral contigs analyzed in this study. Each contig has direct terminal repeats (DTRs) and encodes at least one hallmark gene of Duplodnaviria phages.

##### *Protein function annotation*

`data_file_S5_ORFs.gb`
Multi-GenBank file containing all contigs with their predicted open reading frames (ORFs), including annotations used for downstream protein-level analyses.

`data_file_S6_plm_annotation.zip`
Contains protein function predictions generated with the Virus Protein Function Protein Language Model (VPF-PLM). Includes per-protein `.csv` files with predicted functions and functional probability matrices, as well as `.pkl` embeddings.

`data_file_S7_vibrant_annotation.zip`
VIBRANT-derived annotations for environmental viral contigs. Includes KEGG, Pfam,

VOG, and VPFAM HMM scan results, and summary figures describing genome size, quality, and gene content.

data_file_S8_HK97_hmm_profiles.zip
Hidden Markov Model (HMM) profiles used to detect HK97-fold proteins. Includes Pfam clan CL0373 ("Phage_coat") profiles and extended sets from the efam database.

data_file_S9_hhpred_annotation.zip
Archive of .hhr output files from HHpred, used for remote homology and function inference. These were manually reviewed for key annotations.

##### *Major Capsid Protein Environmental and Evolutionary Analyses*

data_file_S10_environmental_HK97_proteins.faa
FASTA file of amino acid sequences for all HK97-fold major capsid proteins (MCPs) detected in environmental viral genomes.

data_file_S11_sequence_based_trees.zip
IQ-TREE outputs for environmental HK97 MCPs, including maximum-likelihood trees, model statistics, and TreeCluster groupings at varying distance thresholds.

data_file_S12_HK97_reference_identifiers.txt
List of protein and genome identifiers used in the reference HK97-fold dataset, including entries for encapsulins, PICIs, GTAs, and phages.

data_file_S13_reference_HK97_proteins.faa
Reference set of HK97-fold MCPs from NCBI RefSeq genomes, representing isolated and taxonomically characterized phages.

data_file_S14_HK97_protein_structures.zip
Protein structure models used for structural phylogenetics. Includes both experimentally resolved structures (PDB) and AlphaFold-predicted models with confidence assessments.

data_file_S15_HK97_structure_based_tree.zip
Structure-based phylogenetic tree of HK97-fold proteins across viruses, encapsulins, PICIs, GTAs, and viral-like environmental entities.

data_file_S16_encapsulin_enriched_MCPs.zip
Encapsulin-enriched MCPs identified across environmental and RefSeq viral genomes. Includes sequence data and genome-level metadata, including host and genomic architecture.

data_file_S17_tail_proteins_non_encapsulin_enriched_genomes.zip
Annotation summaries for tail-related proteins detected in DTR-containing twilight-zone genomes lacking encapsulin domains in their HK97 MCPs.

### Section 3. Supplementary tables

**Table S1**. Prevalence of contigs across biomes. The table displays the counts for each biome and genome length group, indicating the number of contigs with MCPs.

| Biome | n | Micro | Mini | Control | n with MCP | Micro With MCP | Mini with_MCP | Control with_MCP |
|---|---|---|---|---|---|---|---|---|
| Freshwater | 1107 | 161 | 375 | 571 | 233 | 2 | 31 | 200 |
| Marine | 1596 | 505 | 873 | 218 | 156 | 5 | 75 | 76 |
| Non-marine saline | 182 | 25 | 105 | 52 | 47 | 1 | 28 | 18 |
| Sediment | 92 | 21 | 39 | 32 | 36 | 0 | 16 | 20 |
| Thermal springs | 33 | 9 | 14 | 10 | 4 | 0 | 0 | 4 |
| Bioreactor | 105 | 13 | 45 | 47 | 44 | 0 | 19 | 25 |
| Bioremediation | 8 | 0 | 0 | 8 | 2 | 0 | 0 | 2 |
| Biotransformation | 5 | 1 | 4 | 0 | 1 | 0 | 1 | 0 |
| Built environment | 38 | 2 | 28 | 8 | 7 | 0 | 3 | 4 |
| Food production | 1 | 1 | 0 | 0 | 0 | 0 | 0 | 0 |
| Lab enrichment | 23 | 0 | 21 | 2 | 10 | 0 | 9 | 1 |
| Lab synthesis | 1 | 0 | 1 | 0 | 0 | 0 | 0 | 0 |
| Modeled | 2 | 1 | 1 | 0 | 0 | 0 | 0 | 0 |
| Solid waste | 49 | 4 | 24 | 21 | 11 | 0 | 5 | 6 |
| Wastewater | 288 | 59 | 146 | 83 | 32 | 1 | 11 | 20 |
| Algae | 15 | 2 | 9 | 4 | 3 | 0 | 2 | 1 |
| Fungi | 7 | 0 | 5 | 2 | 1 | 0 | 1 | 0 |
| Human | 2867 | 515 | 1033 | 1319 | 562 | 36 | 212 | 314 |
| Invertebrates | 55 | 12 | 36 | 7 | 14 | 0 | 12 | 2 |
| Mammals | 413 | 24 | 291 | 98 | 24 | 0 | 9 | 15 |
| Microbial | 12 | 3 | 7 | 2 | 0 | 0 | 0 | 0 |
| Plants | 110 | 13 | 30 | 67 | 19 | 0 | 8 | 11 |
| Cave | 7 | 2 | 3 | 2 | 0 | 0 | 0 | 0 |
| Deep subsurface | 54 | 11 | 10 | 33 | 10 | 0 | 4 | 6 |
| Geologic | 2 | 1 | 0 | 1 | 1 | 0 | 0 | 1 |
| Oil reservoir | 2 | 0 | 2 | 0 | 2 | 0 | 2 | 0 |
| Peat | 21 | 4 | 17 | 0 | 1 | 0 | 1 | 0 |
| Plant litter | 10 | 4 | 1 | 5 | 0 | 0 | 0 | 0 |
| Rock-dwelling (endoliths) | 3 | 0 | 2 | 1 | 0 | 0 | 0 | 0 |
| Soil | 349 | 73 | 141 | 135 | 73 | 0 | 29 | 44 |
| Volcanic | 1 | 0 | 1 | 0 | 0 | 0 | 0 | 0 |

**Table S2.** Prevalence of tail genes. The table summarizes the counts of tail genes across the genome length groups (micro, mini, and control), distinguishing those with and without encapsulin enrichment.

| Genome length group | Encapsulin domains presence | n | Genomes with tails count | %with tail | n tails = 1 | n tails >1 | Max tail count | n tail proteins |
|---|---|---|---|---|---|---|---|---|
| mini | FALSE | 398 | 14 | 3.52 | 14 | 0 | 1 | 14 |
| mini | TRUE | 93 | 1 | 1.08 | 1 | 0 | 1 | 1 |
| micro | FALSE | 45 | 8 | 17.78 | 5 | 3 | 4 | 13 |
| micro | TRUE | 0 | 0 | 0 | 0 | 0 | 0 | 0 |
| control | FALSE | 778 | 97 | 12.47 | 81 | 16 | 5 | 124 |
| control | TRUE | 20 | 0 | 0 | 0 | 0 | 0 | 0 |
